# Transient CB2 receptor activation triggers irreversible luminal differentiation via chromatin remodeling in breast cancer

**DOI:** 10.1101/2025.07.29.667375

**Authors:** Nuria G. Martínez-Illescas, Gemma Pérez-García, María Rubert-Hernández, Camila Quezada-Gutiérrez, Estrella Martín-Zapater, Luis M. Alonso-Colmenar, Carmen Hernández, Ángela Zarco-Cuadrillero, Eduardo Caleiras, Miguel Muñoz-Ruiz, Ana Payo-Payo, Laura Frías-Aldeguer, Cristina Sánchez, Mercedes Balcells-Camps, Elazer R. Edelman, María Salazar-Roa

## Abstract

Cellular plasticity enables cancer cells to escape therapy by adopting stem-like or alternate lineage states. Here, we identify a mechanism by which cannabinoid receptor 2 (CB2R) activation promotes irreversible lineage commitment in breast cancer. Using patient-derived and murine organoids, we show that brief, low-dose exposure to CB2R agonists—either phytogenic or synthetic—induces a basal-to-luminal transition, accompanied by reduced self-renewal, invasiveness, and tumor-initiating potential. These changes are retained under conditions that normally promote dedifferentiation, including fibroblast co-culture, immune pressure, and mechanical shear stress.

Mechanistically, CB2R engagement initiates a transient chromatin remodeling program, marked by early expression of pluripotency-associated genes followed by silencing and differentiation commitment. This epigenetically stabilized state renders tumor cells more responsive to tamoxifen and limits the emergence of resistant clones.

Our findings uncover a previously unrecognized role for CB2R in modulating cancer cell identity and suggest new opportunities to constrain tumor plasticity by directing differentiation through a drug-responsive pathway.

## Introduction

Recent evidence of cellular plasticity and spontaneous de-differentiation suggests that cancer developmental landscape is far more complex than a simple hierarchical structure. “Phenotypic plasticity,” a newly recognized hallmark of cancer, plays a significant role in this complexity (1). Traditional cancer treatments, such as cytotoxic drugs and radiotherapy, primarily aim to target cell division through mechanisms of widespread cell damage and toxicity. However, these approaches often prove inefficient and may create evolutionary pressures that favor the emergence of aggressive, resistant cancer clones. In some cases, cytotoxic treatments even promote undesirable side effects, such as metastasis.

In contrast, differentiation therapies aim to induce terminal differentiation in cancer cells, effectively removing them from the proliferative compartment (2, 3). By promoting differentiation in a subset of tumor cells, these therapies can alter tumor dynamics by limiting the growth potential of the entire tumor. This effect is akin to habitat fragmentation, where differentiated cells occupy space and resources that would otherwise be used for tumor expansion, thereby restricting the carrying capacity. Notably, when differentiation strategies are combined with cytotoxic therapies, tumor dynamics can shift from growth to regression (3).

Tumors with high stem cell traits are, therefore, ideal candidates for differentiation-based and potentially, combined therapies. The mammary gland, a tissue characterized by significant postnatal development and cyclical expansion, exhibits considerable stem cell plasticity. This plasticity is particularly relevant in the context of poorly differentiated breast carcinomas, which are known for their rapid spread and high incidence of metastasis, making them a major challenge in cancer treatment. Indeed, a significant proportion of breast cancer-related deaths are attributed to relapse or dissemination after surgery/therapy, a phenomenon most commonly observed in patients with poorly differentiated tumors (4). While the direct relationship between tissue differentiation and partial pathologic complete response (pCR) after treatments remains unclear (5, 6), existing correlations emphasize the need of differentiation-based treatment approaches.

Designing effective differentiation therapies requires an understanding of the parallels between cancer and embryonic development. Many signalling pathways involved in embryogenesis are aberrantly reactivated in cancer, with genetic and epigenetic changes driving this process (7–11). These pathways, including Wnt/β-catenin (12), Hedgehog (Hh) (13), Notch (14), TGF-β (15), PI3K /AKT (16) and JAK/STAT (17), govern cellular differentiation during normal development. In cancer, their reactivation contributes to disease pathology, driving the reprogramming of tumor cells to a progenitor-like state—a key feature of metastasis. Notably, the endocannabinoid system (ECS) has been shown to regulate many of these pathways (18).

The ECS is a complex cellular network that regulates a variety of physiological functions (19, 20). It consists of two main cannabinoid receptors, CB1R and CB2R, their endogenous ligands (AEA and 2-AG), and associated proteins responsible for their synthesis, transport, and degradation. Over the past two decades, substantial evidence has demonstrated that ECS activation has anti-tumor effects in cell and animal models, including promoting tumor cell death, inhibiting endothelial cell migration, blocking tumor angiogenesis, and suppressing tumor cell adhesion and migration (21, 22). Much of this research has focused on cannabinoids derived from *Cannabis sativa*, which contains over 500 compounds, including the primary cannabinoids Δ9-tetrahydrocannabinol (THC) and cannabidiol (CBD) (23). Additionally, synthetic cannabinoid receptor ligands have shown similar effects in cancer models (24). These findings have prompted clinical trials investigating the safety and efficacy of cannabinoids as anti-tumor agents (25–28).

In this study, we expand the therapeutic potential of cannabinoids by exploring their use as differentiation drivers in cancer. Using three-dimensional organoids derived from breast cancer mouse models and patient biopsies, we demonstrate that both natural and synthetic Cannabinoid Receptor 2 (CB2R) ligands promote long-term and stable tumor cell differentiation when administered at low doses (nanomolar concentrations) and short treatment regimens (4 days followed by drug withdrawal). Moreover, combining these differentiation strategies with tamoxifen—a widely used breast cancer treatment—significantly enhances therapeutic outcomes while allowing for reduced chemotherapy doses and exposure, thus mitigating undesirable side effects.

## Results

### Low-dose, transient THC exposure durably reprograms mammary tumor organoids, disrupting cancer cell plasticity *in vitro* and *in vivo*

Cannabinoids have been widely studied for their anti-tumor properties—suppressing proliferation, invasion, metastasis, and promoting autophagy and cell death [recently reviewed in (18)]. However, their role in regulating tumor differentiation remains poorly understood. Since differentiation therapies align with adaptive treatment strategies aimed at reducing tumor evolution and resistance (29), we first examined whether low-dose THC could durably modulate tumor cell plasticity.

To test this, we exposed MMTV-PyMT mammary tumor organoids to nanomolar vs. micromolar concentrations of THC. While high doses triggered expected cytotoxic effects, 10 nM THC induced significant long-term morphological and molecular changes exclusively in 3D cultures, but not in 2D (**Supp. Fig. 1**). Based on this, we developed a protocol using a 4-day 10 nM THC pulse, followed by extended drug-free culture (**Fig. 1A**). Strikingly, this short, low-dose exposure consistently induced a basal-to-luminal switch, both morphologically (**Fig. 1B–E**) and at the molecular level (**Fig. 1F–G**), highlighting a stable reprogramming effect on tumor cell identity. These effects were replicated in organoids derived from MMTV-Her2/neu tumors (**Supp. Fig. 2**), confirming that THC-driven differentiation is not model-specific. Functionally, THC treatment inhibited collective migration, triggered a mesenchymal-to-epithelial transition (MET) (**Fig. 2A–D**), and reduced stem-like features and self-renewal capacity (**Fig. 2E–G**). These phenotypes were also validated in patient-derived breast cancer organoids (**Supp. Fig. 3**), underscoring their potential clinical relevance.

**Figure 1.**
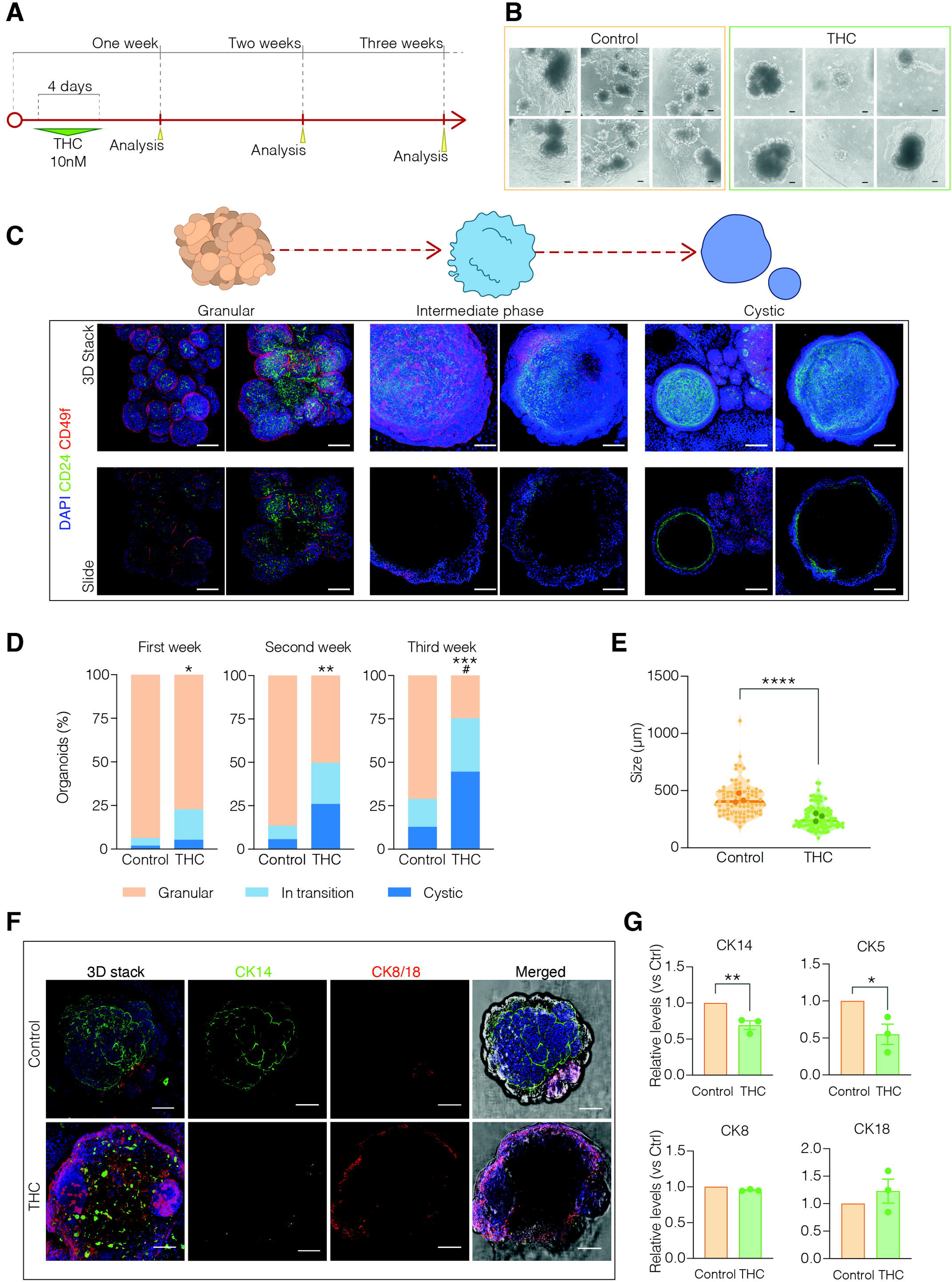
A brief nanomolar THC exposure induces a long-term basal-to-luminal morphological and molecular switch in MMTV-PyMT mammary tumor organoids. **A**, Schematic of the experimental design. PyMT-derived tumor organoids were treated with vehicle (DMSO) or 10 nM THC for 4 days, followed by extended culture. Analyses were performed at 1-, 2-, and 3-weeks post-treatment. **B**, Representative brightfield images of control and THC-treated organoids at 2 weeks. *n*=6 independent experiments. Scale bar, 100 μm. **C**, Confocal images of organoids stained for CD24 (green) and CD49f (red) -cell surface markers of mammary gland precursor-at 1, 2, and 3 weeks, showing temporal changes in morphology and marker expression after THC exposure. DAPI (blue) stains nuclei. Upper panels show 3D stacks; lower panels show representative single slices. *n*=3 independent experiments. Scale bar, 100 μm. **D**, Quantification of organoids classified as granular (basal), mid (intermediate), or cystic (luminal/rounded) over time. *n*=3 independent experiments; *p<0.05, **p<0.01, ***p<0.001 vs. control (granular); ^#^p=0.014 vs. control (cystic) (Two-way ANOVA). **E**, Quantification of organoid size (μm) at endpoint. *n*=3 independent experiments; 80–100 organoids per condition; ****p<0.0001 (unpaired Student’s t-test). **F**, Confocal images of organoids at 2 weeks stained for CK14 (green; basal marker) and CK8/18 (red; luminal marker). Left: 3D stacks; middle: single slices; right: merged with brightfield. DAPI (blue) stains nuclei. *n*=4 independent experiments. Scale bar, 100 μm. **G**, qPCR analysis of cytokeratin genes (CK14/5 and CK8/18, denoting a basal-like and luminal-like phenotypes, respectively) at endpoint, normalized to GAPDH. *n*=3 independent experiments; *p<0.05, **p<0.01 (unpaired Student’s t-test).

**Figure 2.**
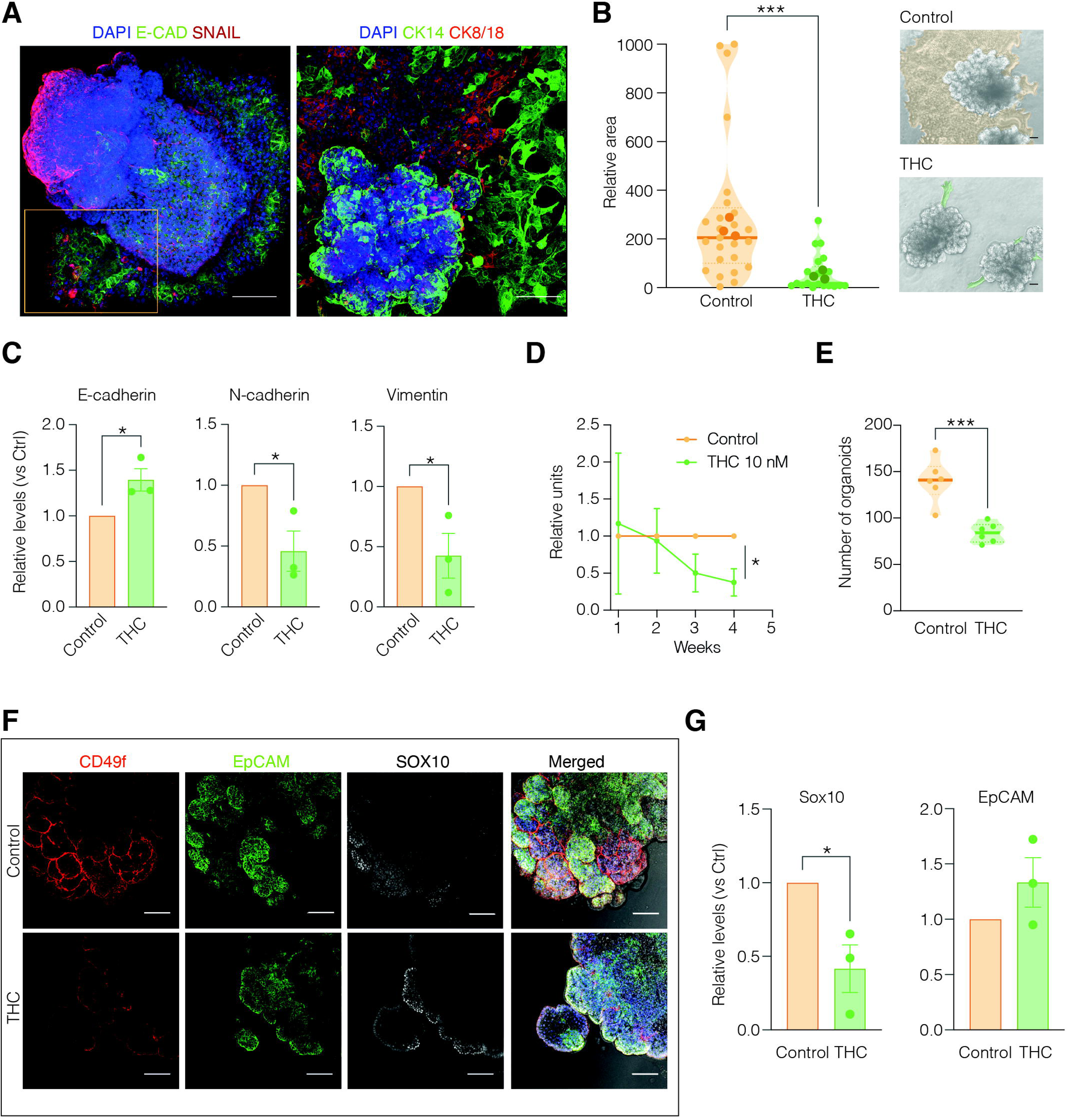
Brief exposure to nanomolar THC prevents collective migration, promotes mesenchymal-to-epithelial transition, and reduces self-renewal in mammary tumor organoids. **A,** Representative confocal images of basal organoids showing collective migration in control cultures. Organoids are stained for E-CAD (green) and SNAIL (red) (left) -as EMT markers- or CK14 (green) and CK8/18 (red) (right). Note the cellular projections extending from the organoid edges and attaching to the substrate. The inset highlights preserved E-CAD staining in migrating areas, indicating cohesive cell movement. DAPI (blue) stains nuclei. *n*=3 independent experiments. Scale bar, 100 μm. **B**, Quantification of the two-dimensional area (μm²) of collective-migratory projections in control vs. THC-treated cultures at 2 weeks. *n*=3 independent experiments; 25–30 organoids per condition; ***p<0.001 (unpaired Student’s t-test). Representative brightfield images (right) illustrate measured areas. Scale bar, 100 μm. **C**, qPCR analysis of EMT-related genes at day 14. mRNA levels were normalized to GAPDH. *n*=3 independent experiments; *p<0.05 (unpaired Student’s t-test). **D**, Time-course comparison of organoids displaying collective migration in control vs. THC-treated conditions. *n*=3 independent experiments; *p<0.05 (unpaired Student’s t-test). **E**, Quantification of secondary organoid formation from single-cell dissociation at 21 days, indicating self-renewal capacity. *n*=6 independent experiments; ***p<0.001 (unpaired Student’s t-test). **F**, Confocal images of organoids stained for CD49f (red), EpCAM (green), and SOX10 (white; stemness marker) after treatment with vehicle or 10 nM THC and analyzed at 2 weeks. DAPI (blue) stains nuclei. Merged images with brightfield are shown on the right. *n*=4 independent experiments. Scale bar, 100 μm. **G**, qPCR analysis of stemness and differentiation markers (Sox10, EpCAM) at day 14. Data normalized to GAPDH. *n*=3 independent experiments; *p<0.05 (unpaired Student’s t-test).

To assess functional *in vivo* consequences, THC-treated and control organoids were orthotopically injected into wild-type PyMT mice (**Fig. 3A–B**). THC-pretreated organoids showed markedly reduced tumor initiation capacity, and the few tumors that did form exhibited delayed growth and non-malignant histology, in contrast to controls which developed aggressive neoplasias (**Fig. 3C–E**). In tail-vein injection assays, THC-pretreated organoids produced significantly fewer lung micro- and macro-metastases compared to controls (**Fig. 3F–I**).

**Figure 3.**
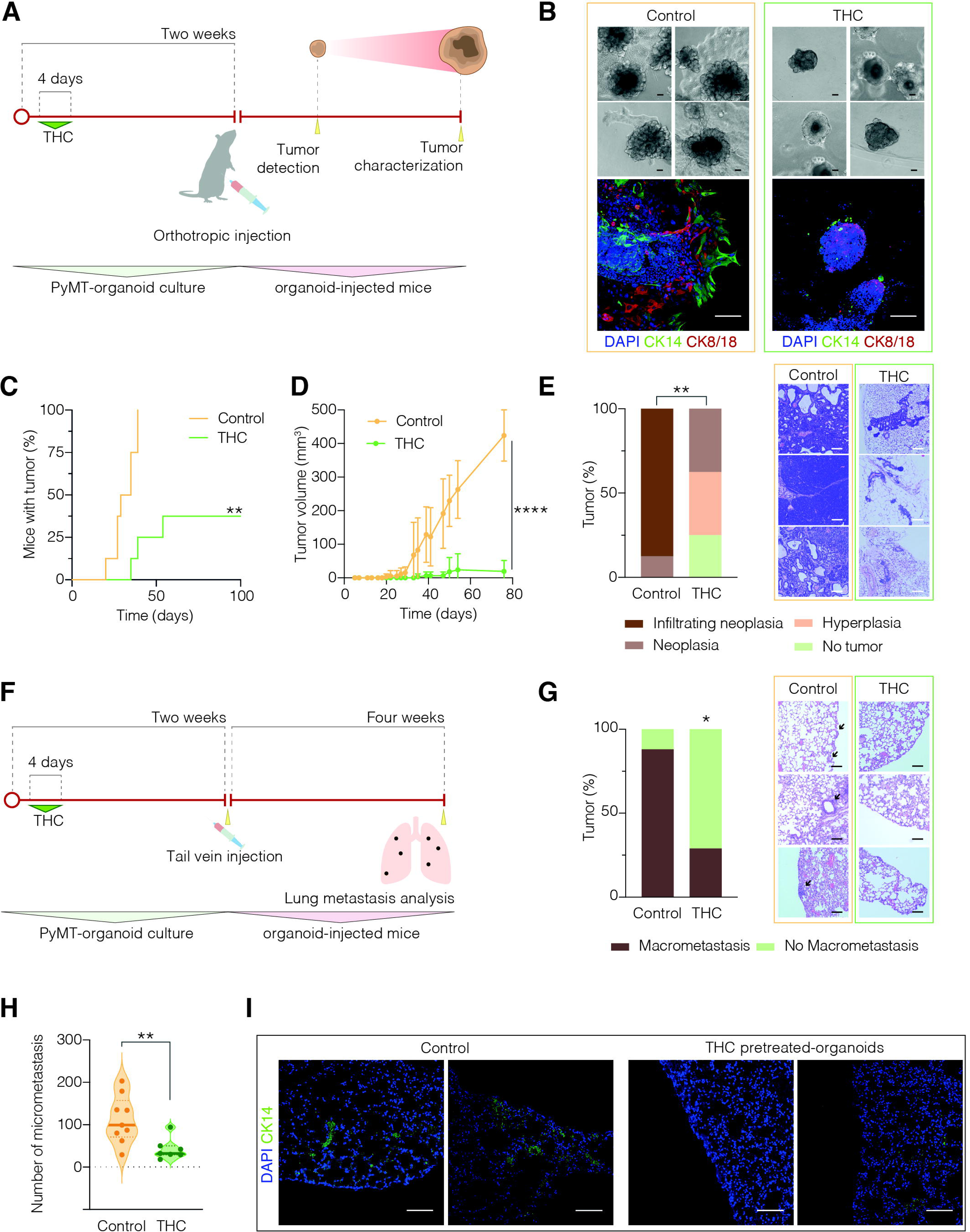
*In vivo* functional evidence of THC-induced tumor cell differentiation. **A**, Schematic of the experimental design. MMTV-PyMT-derived organoids were treated with vehicle or THC (10 nM) for 4 days, cultured drug-free for 10 days, and orthotopically injected into PyMT wild-type littermates. Mice (*n*=8 vehicle, *n*=8 THC) were monitored for 100 days to assess tumor onset and growth. **B**, Brightfield and immunofluorescence images of organoids treated as indicated and analyzed before injection. Organoids showed expected expression of basal (CK14; green) or luminal (CK8/18; red) markers. DAPI (blue) stains nuclei. Scale bar, 100 μm. **C**, Kaplan-Meier plot showing tumor onset (% of mice with tumors) over time. **p<0.01 (Two-way ANOVA). **D**, Quantification of tumor volume (mm³) in both groups. ****p<0.0001 (Two-way ANOVA). **E**, Histopathological classification of tumors at endpoint (100 days): expanded/focal infiltrating neoplasia, local neoplasia, hyperplasia, or no tumor cells. **p<0.01 (Two-way ANOVA). Right: H&E-stained images of representative infiltrating neoplasia (control) and localized hyperplasia (THC). Scale bar, 500 μm. **F**, Schematic of metastasis assay. Organoids were treated with vehicle or THC (10 nM) for 4 days, cultured for 2 additional weeks, and injected via tail vein into PyMT wild-type mice. Lung metastases were assessed one-month post-injection (*n*=9 vehicle, *n*=7 THC). **G**, Distribution of micro- and macro-metastases in lung tissue per condition. *p<0.05 (Two-way ANOVA). Right: H&E-stained images of representative lung metastases. Scale bar, 500 μm. **H**, Quantification of micro-metastases per animal. Each dot represents one mouse. **p<0.01 (Two-way ANOVA). **I**, Immunofluorescence of lung sections stained for CK14, indicating micro-metastases. DAPI (blue) stains nuclei. Scale bar, 500 μm.

Together, these findings reveal that brief, low-dose THC exposure durably reprograms tumor cell fate, reducing plasticity, stemness, and metastatic potential both *in vitro* and *in vivo*.

### CB2 receptor mediates cannabinoid-induced differentiation in breast cancer organoids

To explore the role of cannabinoid receptor tone in tumor cell differentiation, we compared the effects of pure phytocannabinoids (THC, CBD), *Cannabis sativa* extracts, and methanandamide (mAEA)—a metabolically stable analog of the endocannabinoid AEA—on MMTV-PyMT-derived mammary tumor organoids. As shown in **Supplementary Fig. 4**, both extracts and pure cannabinoids induced similar morphological changes, reduced collective migration, and promoted a basal-to-luminal molecular switch. In contrast, mAEA did not equally influence differentiation, maybe due to its low affinity for CB2R. We next assessed synthetic inverse agonists of cannabinoid receptors: SR1 (CB1R) and SR2 (CB2R). Interestingly, SR2, but not SR1, recapitulated THC effects—inducing durable basal-to-luminal reprogramming, inhibiting migration (**Fig. 4A–E**), and reducing self-renewal (**Fig. 4F**). This unexpected finding suggested a functional role for CB2R in regulating tumor plasticity. Combining THC with either SR1 or SR2 did not block its long-term effects on morphology, migration, or organoid size (**Supp. Fig. 5A–D**), implying that CB2R modulation is sufficient but not necessarily required for THC-mediated reprogramming. Supporting this, CB2R expression modestly increased following cannabinoid treatment in differentiation-responsive contexts (**Supp. Fig. 5E**).

**Figure 4.**
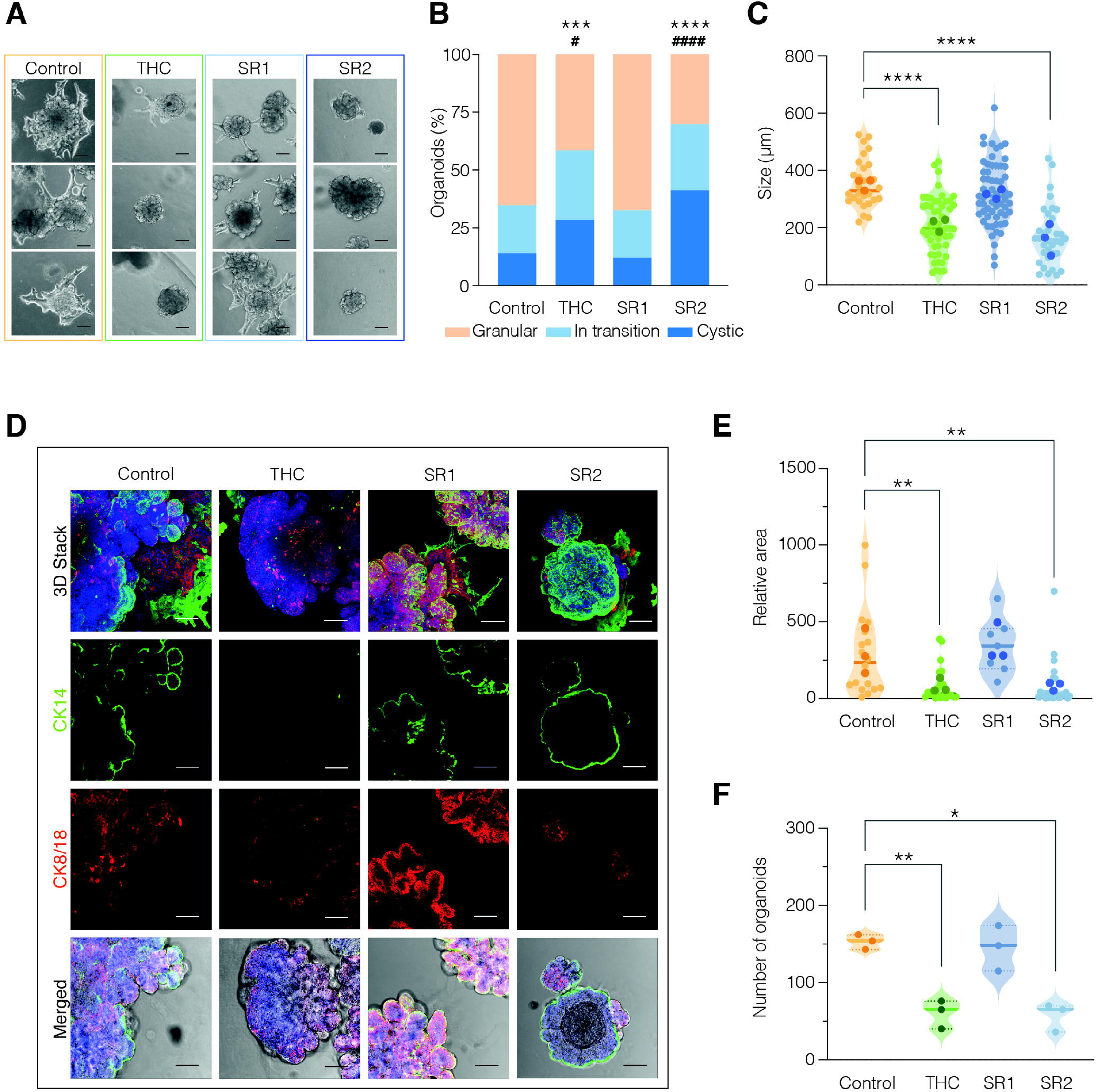
SR2, but not SR1, mimics THC in promoting a basal-to-luminal switch, blocking collective migration, and reducing self-renewal in mammary tumor organoids. **A**, Representative brightfield images of organoids treated for 4 days with vehicle, THC (10 nM), SR1 (100 nM), or SR2 (100 nM), and analyzed after two weeks of drug withdrawal. *n*=4 independent experiments. Scale bar, 100 μm. **B**, Quantification of organoid morphology (granular, mid, cystic) in each condition at 14 days. ***p<0.001; ****p<0.0001 (Two-way ANOVA, granular vs. control); ^#^p<0.05; ^####^p<0.0001 (cystic vs. control). **C**, Organoid size (μm) measured at 14 days. *n*=80–100 organoids/condition across 4 experiments; ****p<0.0001 (One-way ANOVA). **D**, Confocal images of organoids stained for CK14 (green) and CK8/18 (red), showing marker changes 2 weeks after treatment. DAPI (blue) stains nuclei; bottom panels show merged IF and brightfield images. *n*=4. Scale bar, 100 μm. **E**, Quantification of collective-migratory projection area (μm²) in all conditions at 2 weeks. *n*=25–30 organoids/condition; **p<0.01 (unpaired Student’s *t*-test). **F**, Number of secondary organoids formed from single-cell dissociation (indicating self-renewal) at 21 days. *n*=3 independent experiments; *p<0.05; **p<0.01 (One-way ANOVA).

To directly test the role of CB2R, we generated CB2R knockout mice (MMTV-PyMT CB2R⁻/⁻). These mice developed tumors with delayed onset compared to wild-type controls (**Supp. Fig. 6A**), consistent with previous observations in other models (30). CB2R deficiency also altered tumor biology—lower Ki67 and SOX gene expression suggested reduced proliferation and stemness (**Supp. Fig. 6B**). While flow cytometry revealed no major shifts in EpCAM/CD49f-defined subpopulations between CB2R⁻/⁻ and WT tumors (**Supp. Fig. 6C**), the capacity to form viable organoids from luminal progenitors was significantly impaired in CB2R⁻/⁻ mice (**Supp. Fig. 6D**). Additionally, CB2R knockout organoids exhibited distinct basal-like morphologies, reduced size, diminished collective migration, and decreased long-term self-renewal (**Supp. Fig. 6E–J**), closely resembling the phenotype induced by THC and SR2.

Together, these findings identify CB2R signaling as a key modulator of tumor differentiation, capable of driving durable reprogramming in breast cancer organoids.

### THC and SR2 trigger an early, transient epigenetic reprogramming in breast cancer organoids, driving sustained cellular differentiation

Cellular differentiation is tightly linked to epigenetic remodeling, yet the connection between CB2R signaling and epigenetic regulation in cancer remains poorly defined (18). To investigate this relationship, we conducted transcriptomic profiling of mammary tumor organoids treated with THC or SR2, focusing on genes related to epigenetic control and stemness. As shown in **Figure 5**, both treatments triggered a distinct, phase-specific gene expression program. By day 3, organoids displayed a transcriptional signature enriched for embryonic and pluripotency-associated genes (**Fig. 5A–D**), reminiscent of the early reprogramming stages observed in ESCs and iPSCs. This transient increase in plasticity was followed by a clear shift toward a differentiation-related transcriptional profile at later time points (**Fig. 5E**), underscoring a structured reprogramming process.

**Figure 5.**
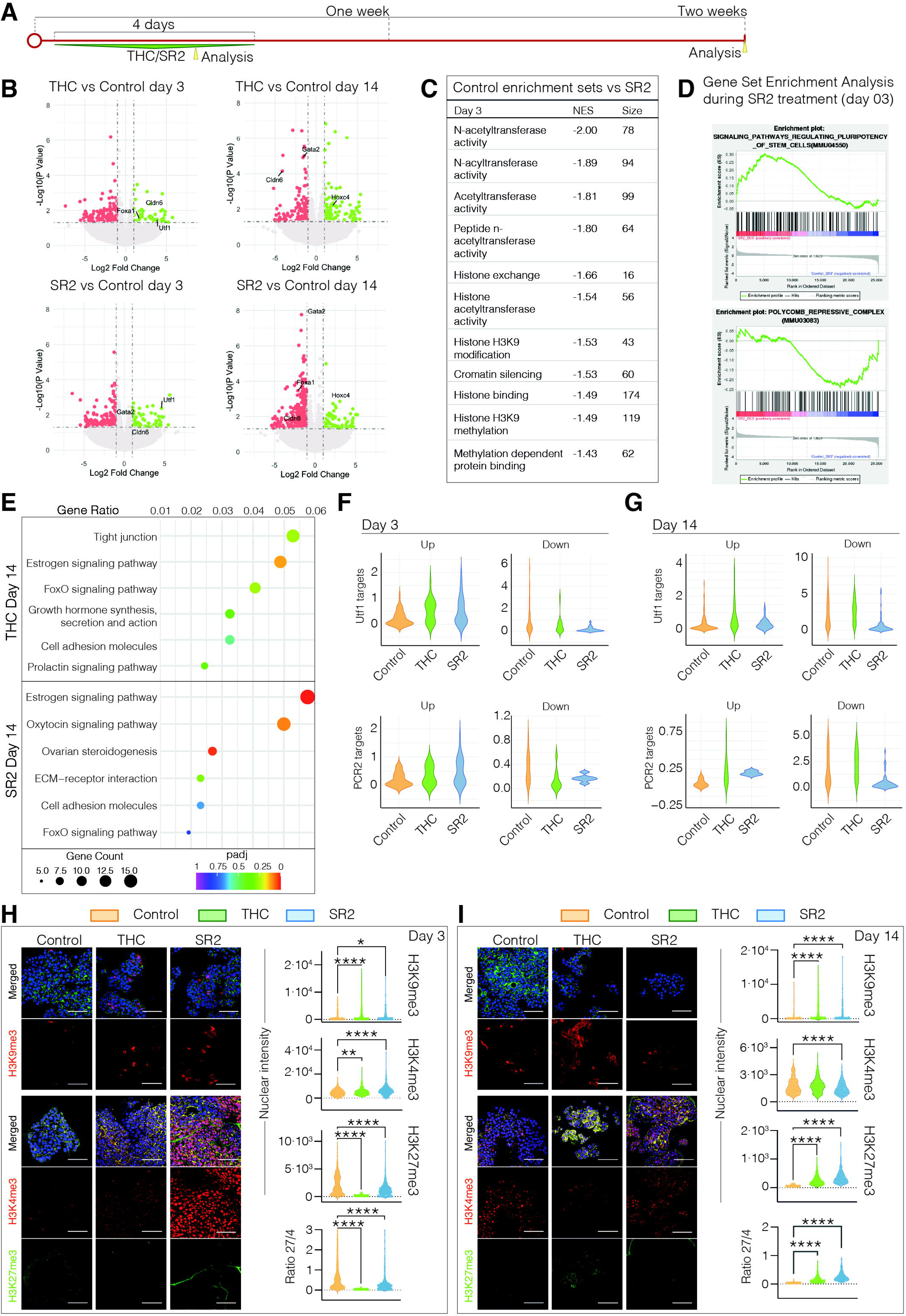
THC and SR2 trigger early and reversible epigenetic remodeling, followed by later differentiation. **A,** Schematic timeline of the experimental workflow. **B**, Volcano plots showing differentially expressed genes in organoids treated with THC (10 nM) or SR2 (100 nM) vs. vehicle at day 3 and day 14 post-treatment. Genes with log₂ fold change between −1 and 1 are considered unchanged. *n*=3 independent experiments. **C,** Gene Ontology (GO) enrichment analysis at day 3 of treatment, showing epigenetic regulation pathways significantly altered in SR2-treated organoids compared to control. Normalized Enrichment Score (NES) and gene set size are indicated. **D,** Gene Set Enrichment Analysis (GSEA) of pathways related to pluripotency (top) and Polycomb Repressive Complex 2 (PRC2; bottom) in SR2-treated vs. control organoids at day 3 of treatment. **E,** Dot plots showing KEGG pathway enrichment analysis of differentially expressed genes following THC (top) or SR2 (bottom) treatment and evaluated at day 14, providing rationale for therapeutic tamoxifen combination in Figure 7. **F,** Violin plots showing expression scores of validated UTF1 (top) and PRC2 (bottom) targets at day 3 (32). **G,** Violin plots of the same gene sets as in (E) at day 14 post-treatment (32). **H-I,** Organoids stained for H3K9me3 (red) and phalloidin (cytoskeleton marker; yellow) in upper images; H3K4me3 (red), H3K27me3 (green) and phalloidin (yellow) in bottom images, at day 3 (**H**) and day 14 (**I**), with DAPI (blue) staining nuclei. Violin plots show quantification of total nuclear fluorescence intensity per nucleus (800-1500 nuclei were evaluated per condition) for every staining, and the H3K27me3/H3K4me3 ratio denotes chromatin bivalency. *n*=3 independent experiments. Scale bar, 100 μm.

Gene set enrichment and differential expression analyses revealed broad modulation of epigenetic regulators involved in histone modifications, chromatin remodeling, and DNA methylation (**Fig. 5C**). These changes aligned with increased accessibility at bivalent chromatin domains (**Fig. 5D, 5F, 5G**), suggesting that CB2R signaling may facilitate transcriptional priming and epigenetic poising of differentiation programs. Indeed, among the top transiently expressed genes, Utf1 emerged as particularly notable—a key regulator of Polycomb Repressive Complex (PRC) activity and pluripotency maintenance (31, 32). Utf1 was upregulated early after THC or SR2 exposure, and subsequently downregulated at later stages, reflecting the transition from a plastic to a more differentiated state (**Fig. 5B**). Consistent with this, the expression of *bona-fide* UTF1 and PRC2 target genes was synchronously modulated under our experimental conditions (**Fig. 5F–G**). Furthermore, we confirmed bivalent histone marks in organoids treated with THC and SR2, which indeed induced a balanced ratio of H3K27me3/ H3K4me3, reinforcing the role of bivalent chromatin regulation in establishing and maintaining cannabinoid-induced differentiation programs (**Fig. 5H–I)**.

Together, these results demonstrate that short-term CB2R activation triggers a transient epigenetic reset, effectively priming mammary tumor organoids for long-term, stable differentiation.

### CB2R-driven differentiation locks tumor cells into a stable, irreversible phenotype

A critical limitation of current differentiation therapies is their potential reversibility, especially under microenvironmental pressures that favor stemness and aggressiveness (2). To challenge the stability of CB2R-induced differentiation, we developed a dynamic co-culture system mimicking a pro-tumorigenic niche. This system included cancer-associated fibroblasts (CAFs), peripheral blood mononuclear cells (PBMCs), and continuous shear stress via a flow-controlled bioreactor (**Fig. 6A and Supp. Fig. 7**). As expected, co-culture with CAFs significantly enhanced tumor cell migration, and reinforced basal and mesenchymal traits in tumor organoids (**Supp. Fig. 7A–E**). These effects, along with a significant increase in organoid size, were further amplified by the inclusion of PBMCs and shear stress (**Supp. Fig. 7F–J**), confirming the model’s capacity to promote cellular plasticity and invasive behavior. Remarkably, even under these challenging microenvironmental conditions, both THC and SR2 treatments effectively reprogrammed tumor organoids toward a luminal, differentiated state and maintained this phenotype over time (**Fig. 6B–E**). Notably, this differentiation state persisted despite prolonged exposure to CAFs, PBMCs, and shear stress, highlighting the durability of cannabinoid-induced reprogramming. To note, cytokine profiling revealed that SR2- and THC-treated co-cultures exhibited a broad activation of immune responses and a reduction in chemokines associated with tumor progression (**Fig. 6F**). Furthermore, cannabinoid-treated organoids displayed sustained epigenetic remodeling, as described above, even within this aggressive microenvironment (**Fig. 6G–H**).

**Figure 6.**
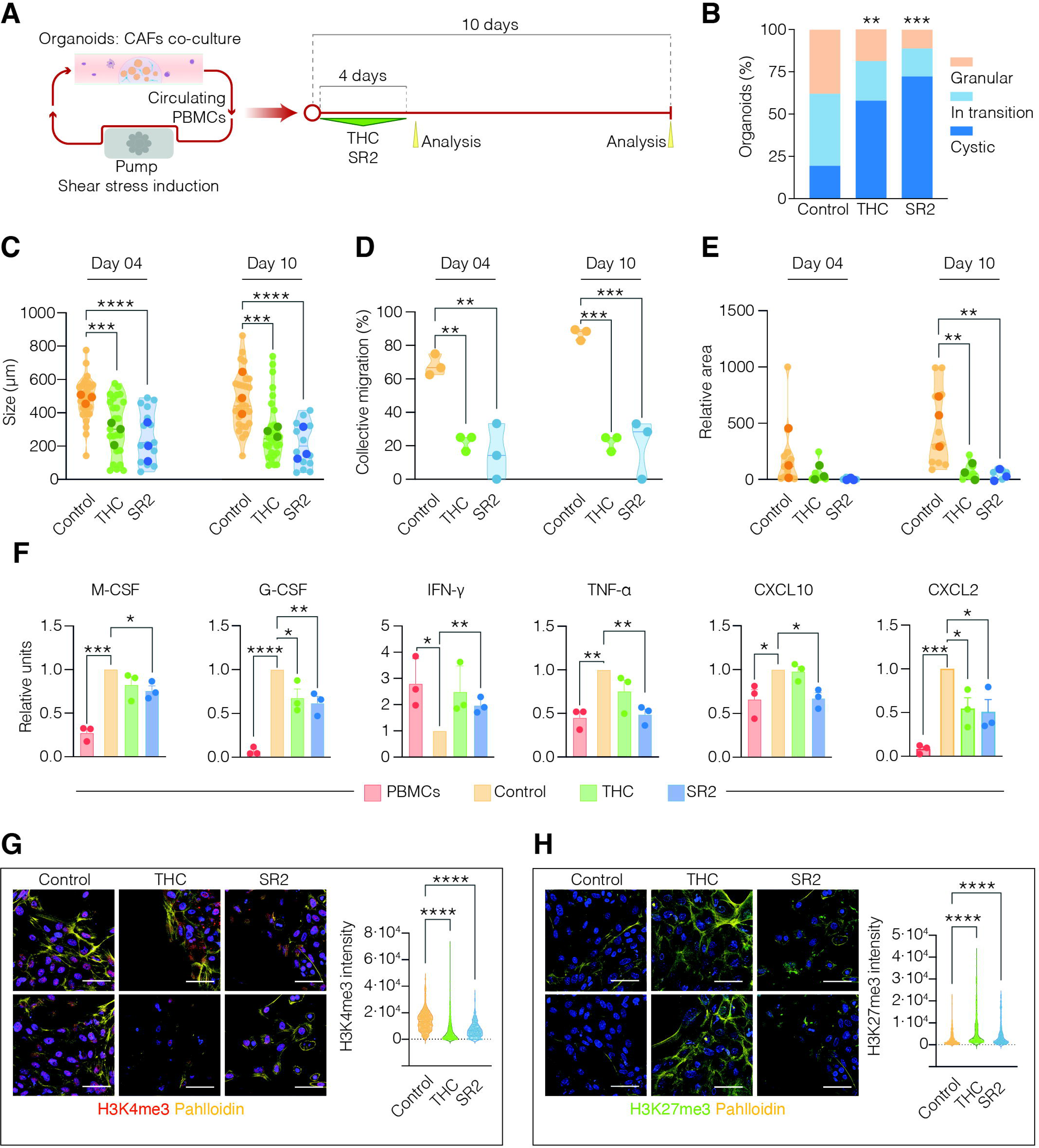
THC and SR2 promote stable differentiation, remodeling organoid architecture, migration, and cytokine signaling in a 3D co-culture system under shear stress. **A,** Schematic of the experimental setup. Organoids embedded in BME were co-cultured with cancer-associated fibroblasts (CAFs) and exposed to circulating peripheral blood mononuclear cells (PBMCs), with continuous pumped flux inducing shear stress. Co-cultures were treated with vehicle, THC (10 nM), or SR2 (100 nM) for 4 days, followed by 10 days in drug-free media. Samples were analyzed at two time points as indicated. **B,** Quantification of organoid morphology (granular, mid, cystic) at day 10 across conditions. *n*=3 independent experiments; *p<0.05 vs. control (granular); ^#^p<0.05 vs. control (cystic) (Two-way ANOVA). **C,** Organoid size (μm) measured at day 4 and day 10. *n*=3 independent experiments; **p<0.01; ***p<0.001; ****p<0.0001 (One-way ANOVA). **D,** Percentage of organoids exhibiting collective migration at day 4 and day 10 in each treatment condition. *n*=3 independent experiments; **p<0.01 (One-way ANOVA). **E,** Quantification of collective migratory projection area (μm²) across conditions at day 4 and day 10. *n*=3 independent experiments; **p<0.01 (One-way ANOVA). **F,** Streptavidin-HRP based analysis of cytokines and chemokines secreted into the culture medium at day 10 for each condition: PBMC alone (red) or organoids+PBMCs+CAFs treated with vehicle (yellow), THC (green) or SR2 (blue). *n*=3 independent experiments; **p<0.01 (One-way ANOVA). **G-H,** Confocal images of organoids at day 10 stained for H3K4me3 (red) and phalloidin (yellow) (**G**) and for H3K27me3 (green) and phalloidin (yellow) (**H**), with DAPI (blue) marking nuclei. Violin plots show the quantification of total nuclear fluorescence intensity per nucleus (800-1500 nuclei were evaluated per condition) for every staining, as indicated. *n*=3 independent experiments. Scale bar, 100 μm.

Collectively, these results demonstrate that CB2R modulation triggers a robust, irreversible differentiation program capable of resisting pro-aggressive cues—supporting its potential as a therapeutic strategy to counteract the inherent plasticity of breast cancer cells.

### CB2R-driven differentiation sensitizes breast cancer cells to endocrine therapy

Drug-resistant cancer cells often incur a fitness cost—reduced growth, higher metabolic demands, or lower competitiveness. Adaptive therapy exploits this trade-off by maintaining a population of drug-sensitive cells to suppress resistant clones through natural competition, thus delaying resistance and improving long-term outcomes (33). In this framework, differentiation therapies, including CB2R-targeted strategies, may act as adaptive interventions by reducing tumor stemness and plasticity, thereby lowering resistance potential. To explore this therapeutic synergy, we tested CB2R-mediated differentiation followed by tamoxifen, a widely prescribed endocrine therapy for ER⁺ breast cancer (34) (**Fig. 7A**). Transcriptomic profiling revealed that both THC and SR2 treatments significantly enhanced estrogen signaling pathways (**Fig. 5E**), providing a rationale for their combination with tamoxifen. Organoids were first exposed to CB2R ligands for 4 days, and ten days after drug withdrawal, treated with sub-micromolar concentrations of tamoxifen for 4 days. This sequential combination therapy led to reduced cell viability (**Fig. 7B**) and G2-M cell cycle arrest (**Fig. 7C**) in cannabinoid-pretreated organoids. After 4 weeks of drug withdrawal, these first observations translated into a more pronounced basal-to-luminal shift (**Fig. 7D**), smaller organoid size (**Fig. 7E**), impaired collective migration (**Fig. 7F**) and particularly relevant, reduced long-term self-renewal (**Fig. 7G**). Importantly, CB2R-based pretreatment completely abolished the emergence of resistant clones, which were evident in tamoxifen-only cultures (**Fig. 7G**). This demonstrates that cannabinoid-induced differentiation not only sensitizes tumor cells to endocrine therapy but also prevents adaptive resistance.

**Figure 7.**
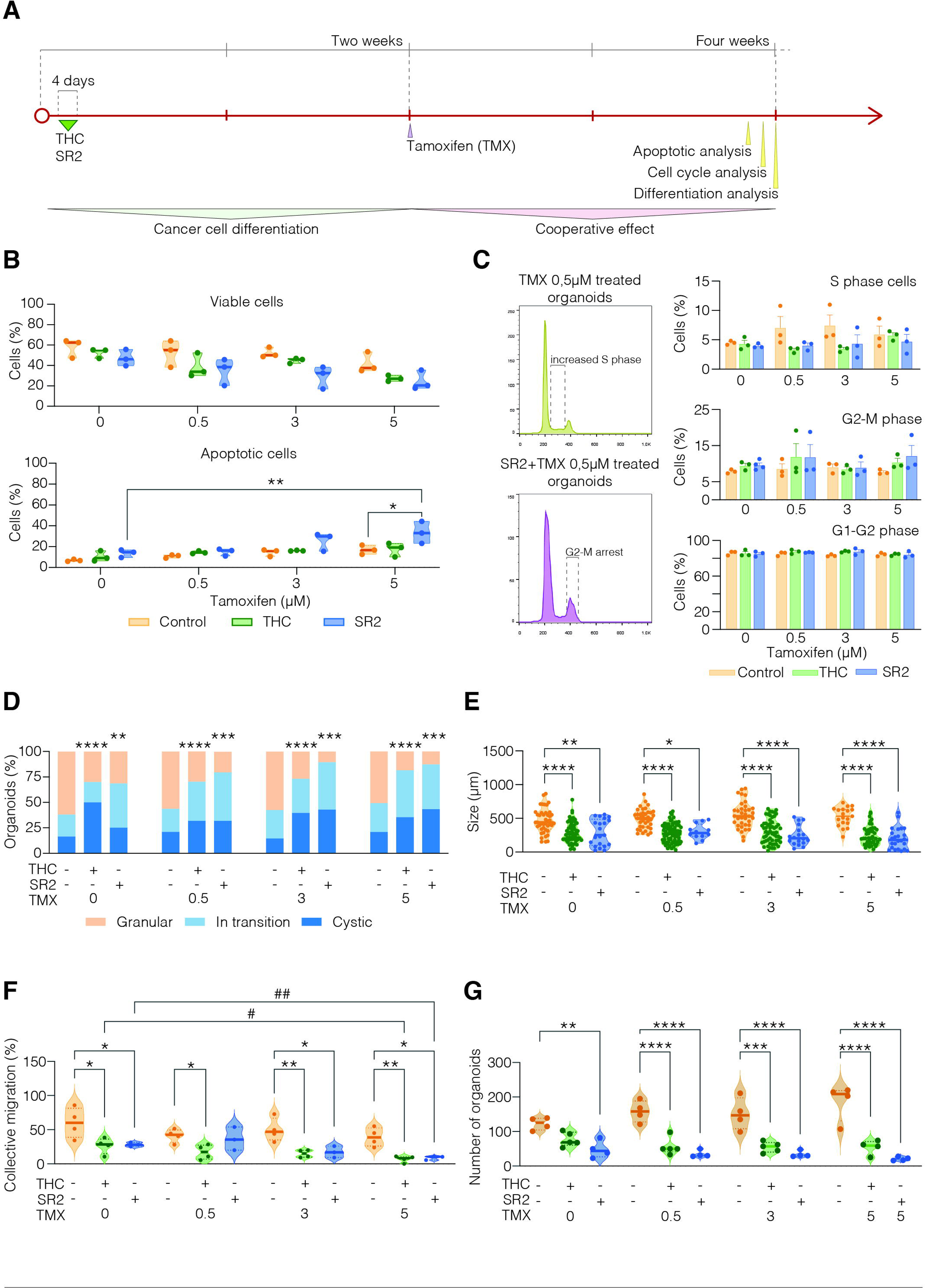
Cannabinoid-induced differentiation enhances tamoxifen response by promoting basal-to-luminal switching and reducing tumor organoid fitness. **A,** Schematic of the experimental design. MMTV-PyMT–derived organoids were pre-treated with vehicle, THC (10 nM), or SR2 (100 nM) for 4 days, followed by 10 days in drug-free media. Tamoxifen (0.5 μM) was then administered for 4 days, and cultures were analyzed immediately or up to 2 weeks later. **B,** Flow cytometry analysis of apoptosis using PI and Annexin V, assessed immediately after tamoxifen exposure or 2 weeks later in pre-differentiated cultures. *n*=4 independent experiments; *p<0.05; **p<0.01; (Unpaired Student’s t-test). **C,** Cell cycle profiles comparing tamoxifen alone to SR2 + tamoxifen at increasing doses (0–5 μM). Left: representative flow cytometry plots. Right: quantification of cell cycle phases (G1, S, G2/M). *n*=3 independent experiments. **D,** Quantification of organoid morphology at endpoint: granular (basal), mid-transition, or cystic (luminal). *n*=4 independent experiments; **p<0.01; ***p<0.001 ****p<0.0001 (Two-way ANOVA). **E,** Organoid size (μm) at endpoint across treatment conditions. *n*=4; 80–100 organoids per condition; *p<0.05; **p<0.01; ****p<0.0001 (One-way ANOVA). **F,** Percentage of organoids exhibiting collective-migratory projections at endpoint. *n*=4; *p-*values as indicated (One-way ANOVA). **G,** Self-renewal capacity assessed by quantification of secondary organoids formed after single-cell dissociation. *n*=4 independent experiments; **p<0.01; ***p<0.001; ****p<0.0001 (One-way ANOVA).

To validate these observations in clinically relevant settings, we tested the combinatory treatment in patient-derived samples. Surgical specimens from two breast tumors were processed for organoid culture, and the efficacy of the SR2–tamoxifen regimen was assessed following the experimental timeline outlined in **Fig. 8A**. Using phase holographic imaging, we analyzed cancer cell movement and dispersal 10 days after CB2R-based differentiation. Cells pretreated with SR2 exhibited reduced motility, altered migration directionality, and changes in cell area and shape irregularity, suggesting impaired adaptation for migration (**Fig. 8B–F and Supp. Fig. 8**). At this same time point, SR2 treatment had also enhanced cell–cell adhesion within the 3D organoids, as evidenced by increased E-cadherin and decreased Snail expression (**Fig. 8G**), along with a marked reduction in organoid size (**Fig. 8H**) and diminished collective migration (**Fig. 8I**). Following this, organoid cultures were exposed to 5 μM tamoxifen for 4 days. Notably, the differentiated cellular state induced by SR2 created a highly favorable context for tamoxifen therapy: organoids pretreated with SR2 showed severely impaired self-renewal following tamoxifen exposure, whereas control organoids retained normal self-renewal capacity despite tamoxifen treatment (**Fig. 8J–K**).Together, these results highlight the potential of CB2R modulation as a priming strategy to enhance the efficacy of standard treatments, aligning with adaptive therapy principles to improve durable cancer control.

**Figure 8.**
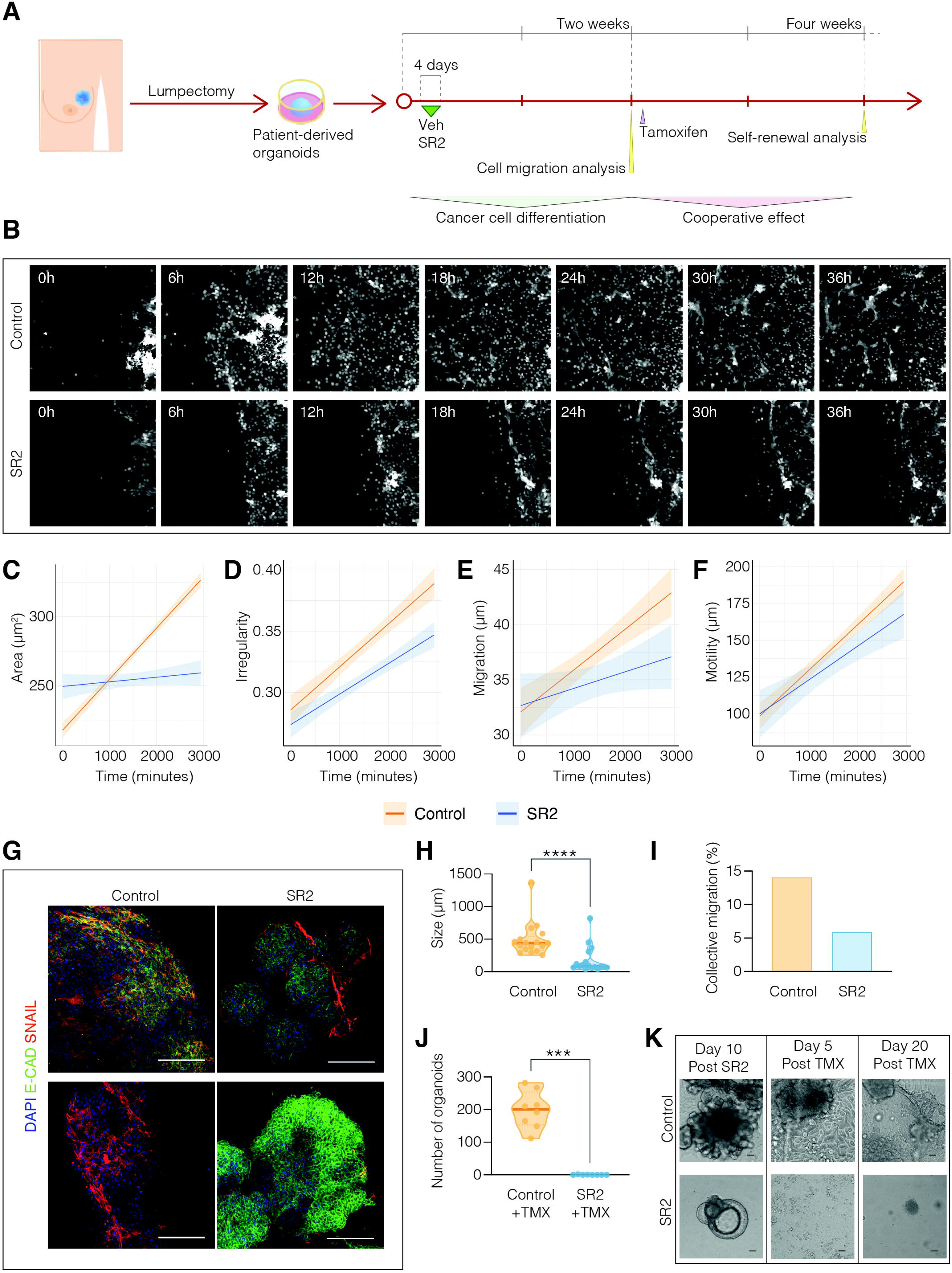
CB2R–tamoxifen combination enhances therapy efficacy in patient-derived breast cancer organoids. **A**, Experimental workflow. Surgical specimens were used to establish patient-derived organoid cultures. After expansion, organoids were treated with vehicle or SR2 (100 nM) for 4 days, followed by a 10-day drug-free period. Prior to tamoxifen treatment, cell dispersal from organoids was assessed using dynamic holographic imaging. Tamoxifen (5 μM) was then administered for 4 days. Organoids were analyzed two weeks later to evaluate cooperative effects on self-renewal. *n*=2 independent tumors, 3 technical replicates each. **B**, Representative holographic images showing 2D movement of organoid-derived cells treated with vehicle or SR2 (100 nM), assessed in a wound healing assay at different time points. **C–F**, Quantification of wound healing assay parameters: cell area (μm²) (**C**), irregularity (**D**), migration distance (μm) (**E**), and motility (μm) (**F**) under both treatment conditions. A linear model was fitted to assess the impact of experimental conditions on each variable. Statistics shown in **Supp. Figure 8**. **G**, Confocal images of organoids stained for E-cadherin (green) and Snail (red), analyzed 10 days post-differentiation. Nuclei were stained with DAPI (blue). Shown are representative 3D confocal stacks. Scale bar = 100 μm. **H**, Quantification of organoid size (μm) at day 10 post-differentiation. ****p < 0.0001 (one-way ANOVA). **I**, Percentage of organoids displaying collective migration at day 10 post-differentiation. **J**, Total number of secondary organoids formed after single-cell disaggregation, reflecting self-renewal capacity, measured two weeks after tamoxifen treatment. ***p < 0.001 (one-way ANOVA). **K**, Representative brightfield images of organoids at different time points: 10 days after differentiation, and 5 or 20 days after tamoxifen treatment, following the experimental timeline shown in (A).

## Discussion

Cancer cells undergo numerous molecular and behavioral changes that enable them to overcome natural selection barriers, one of the most notable being the loss of differentiation markers and a reprogramming to a de-differentiated state. This reversion to an earlier developmental stage provides the tumor with increased plasticity and self-renewal capacity.

The standard treatment for cancer traditionally relies on cytotoxic drugs and radiotherapy, both of which induce cell damage and death, especially in highly proliferative cells. However, these therapies are often criticized for their side effects and their tendency to trigger evolutionary pressures, leading to the emergence of resistant and aggressive tumor clones. As a result, there is growing interest in integrating additional therapeutic approaches, including personalized medicine and immune-directed therapies, to improve clinical outcomes. Among the most promising alternatives to cytotoxic treatments are differentiation therapies, which, although proposed over 50 years ago (35) are still being explored. The principle behind these therapies is that cancer cells, to expand and maintain plasticity, often block differentiation processes. Differentiation therapy does not aim to induce cell death but instead focuses on inhibiting self-renewal and guiding tumor cells back toward terminal differentiation.

One of the most successful examples of differentiation therapy is the treatment of Acute Promyelocytic Leukaemia (APL), where a combination of cytotoxic therapy and all-trans retinoic acid (RA) has dramatically improved remission rates and survival outcomes (36, 37). Notably, while RA alone induces high differentiation grades, long-term disease remission is not achieved, suggesting that combination therapies yield the best results. In solid tumors, differentiation therapy is still in its early stages but remains a promising second-line option for patients with advanced cancer.

Our study uncovers a previously unrecognized role for the cannabinoid receptor CB2 (CB2R) in driving stable cancer cell differentiation in breast cancer. We show that a brief, low-dose exposure to THC or the CB2R ligand SR2 durably reprograms mammary tumor organoids, triggering a basal-to-luminal switch, reducing stemness, and impairing migration, self-renewal, and metastatic potential—effects that persist *in vitro* and *in vivo*.

Over the past decades, substantial preclinical and clinical evidence has demonstrated that the endocannabinoid system (ECS) regulates several physiological functions, including pain perception, motor coordination, and nausea/vomiting (38), common side effects of chemotherapy. Our lab was among the pioneers in documenting the anti-tumor properties of cannabinoids (39), showing their ability to block proliferation, migration, and angiogenesis (22). These findings laid the groundwork for the first phase 1b randomized, placebo-controlled clinical trial assessing the effects of the cannabis-based drug Sativex on tumor progression in patients with recurrent glioblastoma. This trial demonstrated the safety, tolerability, and lack of interaction with temozolomide (the standard treatment), as well as a notable survival benefit at one year (27). With the increasing legalization of cannabis in over 80 countries, there has been a growing interest in its clinical applications, particularly in cancer treatment.

Our study contributes to the expanding body of research on cannabinoids in cancer therapy by focusing on a relatively unexplored application: While cannabinoids have been implicated in nearly all cancer hallmarks (40), their role in differentiation has received less attention. Previous studies, particularly in leukaemia, have shown that THC can sensitize cancer cells to chemotherapeutic agents, reducing IC50 values by approximately 50% and enhancing overall therapeutic effects (41). Additionally, Dronabinol, an FDA-approved cannabinoid, has been shown to inhibit differentiation blockage in acute leukaemia cells (42). In myelomas, THC has been found to induce differentiation of myeloid-derived suppressor cells, reducing immune evasion (43). Most studies have used cannabinoid concentrations in the micromolar range (1-25 μM) (44, 45), with effects typically observed after prolonged exposure. In contrast, our study is the first to evaluate the anti-cancer effects of cannabinoids at nanomolar concentrations (10 nM) in three-dimensional organoid models derived from both mouse and human solid tumors. Moreover, we assess the long-term impact of a brief cannabinoid exposure (4 days), followed by drug withdrawal for at least two weeks, a novel approach that has not been previously explored. Significantly, this work also shows that, after cannabinoid treatment, tumors can be subjected to chemotherapy at reduced doses, resulting in improved therapeutic outcomes. Therefore, our data illustrate a new potential practice of adaptive therapy, with important implications in tumor cell plasticity and relapse impairment.

Intriguingly, we found that THC, SR2, and CB2 receptor knockout (CB_2_KO) cells exhibit similar effects on cancer cell differentiation. Although THC is known to be a partial agonist at both CB1 and CB2 receptors (46) and SR2 is an inverse agonist of CB2R, THC may act also as an inverse agonist at CB2 receptors in a stemness context, or its brief and low-dose exposure might stimulate a CB2-CB1 receptor switch, similar to what has been observed in neuronal differentiation (47–49). This hypothesis aligns with the idea that CB2 receptors are overexpressed in cancer, contributing to tumor aggressiveness, and that cancer differentiation would involve the downregulation of CB2 receptors and a (potential) subsequent upregulation of CB1 receptors (50). Those exciting possibilities are out of the scope of this work but deserve broader research.

Our work shows that brief exposure to CB2R ligands such as THC or SR2 significantly enhances the long-term differentiation potential of cancer cells, even after multiple passages, suggesting an epigenetic mechanism. In fact, RNAseq analyses revealed and early remodeling of the epigenetic landscape, apparently establishing a form of replication memory, that promoted a sustained differentiated state several weeks after treatment. Among the top genes regulated by THC or SR2, Undifferentiated Transcription Factor 1 (UTF1) likely plays an important role in this effect. Extensive research supports its function as a central mediator of epigenetic stability, both in early embryogenesis (51), governing lineage specification, and in cancer, when it has been showed to bind chromatin with histone-like dynamics, maintaining nuclear structure during cell fate transitions (31, 52, 53). Mechanistically, UTF1 localizes to promoters of developmental genes—particularly those that are silenced or bivalently marked (e.g., H3K4me3 and H3K27me3), fine tuning the balance between self-renewal and differentiation (32). Indeed, its expression diminishes as cells begin to differentiate. Collectively, these findings position UTF1 potentially explaining the long-term effects of transient CB2R ligand exposure on cancer cell differentiation.

Synthetic CB2R ligands have shown great promise in pharmacological research, with conceivable applications in neurodegenerative diseases, autoimmune disorders, chronic pain, and cancer (54). However, clinical testing of CB2R-targeted compounds is still in its early stages. Our findings provide strong support for the therapeutic potential of SR2 as differentiation agent in cancer, since our data show that SR2-treated organoids exhibit even stronger differentiation effects than those treated with THC. These results, combined with recent advances in CB2R pharmacology, suggest a promising future for CB2R synthetic ligands in novel cancer therapy.

## Methods

### Animal models and procedures

All animal procedures were approved by the Animal Experimentation Committee of the Complutense University of Madrid (CEIyBA) and the Regional Government of Madrid, in compliance with European animal research regulations. Mice were housed under a 12-hour light/dark cycle with ad libitum access to food and water. PyMT mice [FVB/N-Tg(MMTV-PyMT)^634Mul/J^] were generously provided by Miguel Quintela (CNIO, Spain). The MMTV-neu:CB2R^-/-^ strain was previously developed in our laboratory (30) and MMTV-neu [FVB/N-Tg(MMTV-neu)^202Mul/J^] mice were obtained by selecting CB2R^+/+^ individuals from this colony. To generate the PyMT:CB2R^-/-^ line, PyMT mice were crossed with MMTV-neu:CB2R^-/-^ mice. The CB2R^-/-^ allele was introduced through four generations of backcrossing, during which the neu transgene was also eliminated. Genotyping was performed using the following primers (5′−3′; F: forward, R: reverse). **PyMT**, F: TGC CGG GAA CGT TTT ATT AG; R: GGA ACA AGT ACC CCA GCT CA; **CB2R-WT/KO**, F: CTA CAA AGC TCT AGT CAC CCG; WT-R: GGC TCC TAG GTG GTT TTC ACA TCA GCC TCT; KO-R: CTA AAG CGC ATG CTC CAG AC; **Neu**, F: GCT CAG AGA CCT GCT TTG GA; R: AGG AGG ACG AGT CCT TGT AGT G. PCR was performed using the following protocol: 95 °C for 15 min; 30 cycles of 94 °C for 30 s, 63 °C for 45 s, 72 °C for 1 min; final extension at 72 °C for 10 min; hold at 4 °C.

### Reagents

Cannabinoid-based treatments included Δ⁹-tetrahydrocannabinol (THC, THC Pharm), cannabidiol (CBD), methanandamide (mAEA, Cayman Chemical, #90050), a high-THC cannabis extract, and a balanced THC:CBD (1:1) extract (kind gift from Dr. Paola Pineda, Grupo Curativa, Colombia). Selective cannabinoid receptor inverse agonists included SR141716A (CB1R, SR1, Tocris Bioscience, #0923) and SR144528 (CB2R, SR2, Tocris Bioscience, #1464). Tamoxifen (Sigma, T5648) was used as a reference endocrine therapy. Stock solutions were prepared for a final 1:1000 dilution. mAEA was dissolved in ethanol; all other compounds (THC, CBD, extracts, SR1, SR2, and tamoxifen) were dissolved in DMSO. Vehicle controls were included in all experiments using matched volumes of the respective solvents (DMSO or ethanol) in the culture medium.

### Primary tumor cell isolation and culture

PyMT mice were euthanized at 6–8 weeks of age upon development of palpable tumors. Tumors were excised and placed on ice in tumor cell medium consisting of DMEM (Corning, 10-017-CV) supplemented with 1× GlutaMAX (Invitrogen, 12634-034), 10% FBS (Gibco, A5256701), and antibiotics. Under aseptic conditions, tumors were minced into small fragments using a sterile scalpel. Minced tissue was incubated in 5 mL of freshly prepared digestion solution (tumor medium supplemented with 0.125 mg/mL Collagenase [Sigma, C9407] and 0.125 mg/mL Dispase II [Gibco, 17105041]), prewarmed to 37 °C. The suspension was digested at 37 °C with agitation (150 rpm) for 2 hours. Post-digestion, the mixture was filtered through a 100 μm cell strainer to remove debris, centrifuged at 400 x g, and resuspended in fresh tumor medium. Cells were plated and incubated at 37 °C for 20 minutes to promote adherence. Non-adherent cells were removed by washing to eliminate fibroblasts and stromal contaminants. Fresh medium was then added. Primary tumor cells were confirmed as epithelial by positive staining for pan-cytokeratin (Abcam, ab7753). Cultures were maintained at 37 °C in a humidified incubator with 5% CO₂, tested weekly for mycoplasma contamination, and passaged every 4 days upon reaching confluence.

### Generation and culture of mammary tumor organoids

PyMT, PyMT:CB2R⁻/⁻, and MMTV-neu mice were euthanized at 6–8 weeks of age, and tumors were harvested for organoid generation. Organoids were established and maintained as previously described (5). Briefly, excised tumors were minced and digested in pre-warmed digestion solution consisting of DMEM supplemented with 1× GlutaMAX, 1% FBS, antibiotics (referred to as wash medium), 0.125 mg/mL collagenase, and 0.125 mg/mL Dispase II. The tissue suspension was incubated at 37 °C with constant shaking (150 rpm) for 2 hours. Following enzymatic digestion, the cell suspension underwent sequential centrifugation at 500 × g to wash away adipose tissue remnants using cold wash medium. The resulting pellet was resuspended in basal medium composed of Advanced DMEM/F-12 (Invitrogen, 12634-034), 1× GlutaMAX, 10 mM HEPES (Invitrogen, 15630-056), and antibiotics, then centrifuged again at 500 × g. Pellets were carefully resuspended in 10 mg/mL cold Cultrex growth factor-reduced Basement Membrane Extract (BME) Type 2 (Trevigen, 3533-010-02), and 40 μL domes were plated in pre-warmed 24-well plates. After the BME solidified, 650–1000 μL of pre-warmed breast cancer (BC) organoid expansion medium was added to each well. The expansion medium was prepared by supplementing the basal medium with the following components: 10% homemade R-Spondin-1 conditioned medium, 1× B27 supplement (Gibco, 17504-044), 1× N2 supplement (Gibco, 17-502-, 10 mM nicotinamide (Sigma, N0636), 1.25 mM N-acetylcysteine (Sigma, A9165), 10 nM Gastrin I (Sigma, G9145), 100 ng/mL FGF10 (Peprotech, 100-26), 50 ng/mL Noggin (Peprotech, 120-10C), 50 ng/mL EGF (Peprotech, AF-100-15), 5 nM Neuregulin-1 (Peprotech, 100-03), 5 μM Y-27632 (Abmole), 500 nM A83-01 (Tocris, 2939), 500 nM SB202190 (Sigma, S7067), and Primocin (Invitrogen, Ant-pm-1). Organoids were passaged weekly, and expansion medium was refreshed every two days. For cannabinoid treatment, organoids were exposed for four days to compounds dissolved in DMSO at the following concentrations: 1 nM to 5 μM Δ9-THC, 10 nM CBD, 10 nM THC-rich plant extract, 5 nM:5 nM THC:CBD 1:1 plant extract, 100 nM SR1, 100 nM SR2, and 10 nM methanandamide (mAEA). Unless otherwise specified in the figure legends or text, all analyses were conducted 10 days after treatment withdrawal.

### Generation and culture of breast cancer patient-derived organoids

Patient-derived tumor samples were minced and digested as previously described (5), using a pre-warmed digestion solution at 37 °C with constant shaking (150 rpm) for 2 hours. The resulting suspension underwent sequential centrifugation at 500 × g. Cells were then carefully resuspended in BME, and 20 μL drops were plated in pre-warmed 24-well plates. After the Matrigel solidified, patient breast cancer (BC) organoid expansion medium was added. The patient BC organoid expansion medium consisted of basal medium supplemented with 20% homemade R-Spondin-1 conditioned medium, 1× B27 supplement, 1× N2 supplement, 10 mM nicotinamide, 1.25 mM N-acetylcysteine, 10 nM Gastrin I, 100 ng/mL FGF10, 100 ng/mL Noggin, 50 ng/mL EGF, 5 nM Neuregulin 1, 5 μM Y-27632, 500 nM A83-01, 500 nM SB202190, and Primocin. Organoids were passaged every 7-10 days, and the BC organoid expansion medium was refreshed every two days. Cannabinoid treatments, dissolved in DMSO, were applied to organoids for four days.

### Cancer-Associated Fibroblast (CAF) extraction and culture

Mammary gland tumors were harvested from PyMT mice at 6–8 weeks of age for CAF isolation, as previously described (55, 56). Tumors were minced into small fragments and digested in a prewarmed solution containing wash medium supplemented with 0.125 mg/mL collagenase, 0.125 mg/mL dispase II, and 0.1 mg/mL DNase I. The suspension was incubated at 37 °C with constant shaking (120 rpm) for 2 hours. Following digestion, the cell suspension was filtered through a 100 μm strainer and centrifuged at 500 x g. The pellet was resuspended in selection medium (DMEM with 10% FBS, 1× Insulin-Transferrin-Selenium [ITS-G; Gibco, 41400045], and antibiotics) and incubated for 30 minutes to allow selective fibroblast adherence. After incubation, the medium was replaced with CAF culture medium (DMEM supplemented with 10% FBS, 1× GlutaMAX, and antibiotics). CAF identity was confirmed by immunostaining: cells were vimentin-positive (BD Biosciences, 5505013) and cytokeratin-negative (Abcam, ab7753).

### Peripheral Blood Mononuclear Cell (PBMC) isolation

Peripheral blood was collected via cardiac puncture from adult PyMT tumor-bearing mice and transferred to EDTA-coated tubes to prevent coagulation. Samples were diluted 1:2 in HBSS (Gibco, 14025092). PBMCs were isolated by density gradient centrifugation using Ficoll-Paque™ PLUS (VWR, 17-1440-02), at a ratio of 1 mL Ficoll per 3 mL of diluted blood. The blood was carefully layered over Ficoll and centrifuged at 1600 rpm for 27 minutes at room temperature (no brake, low acceleration). The PBMC layer was collected, transferred to a clean tube, and washed three times with PBS supplemented with 1% FBS and antibiotics. Cells were counted and cryopreserved for subsequent co-culture experiments.

### Mammary tumor organoid–CAF co-culture

PyMT-derived organoids were dissociated and co-seeded with primary CAFs at a 1:250 organoid-to-CAF ratio. CAFs were harvested from confluent cultures by incubation with 0.25% trypsin and 0.1% EDTA (Corning, 25-053-CI) for 4 minutes at 37 °C. Detached cells were collected in CAF medium, centrifuged at 600 x g, and kept on ice until use. Simultaneously, organoids were harvested by mechanical disruption and washed twice by centrifugation at 500 x g. For co-culture, 250 CAFs per organoid were added to the organoid suspension, followed by centrifugation at 500 x g. The resulting pellet was gently resuspended in cold BME, and 20 μL drops were plated into prewarmed 8-well chamber slides (Ibidi, 80841). After BME polymerization at 37 °C, 200–300 μL of prewarmed breast cancer organoid expansion medium was added per well. As experimental controls, organoids were cultured under identical conditions either alone (without fibroblasts) or co-seeded with NIH/3T3 fibroblasts (ATCC CRL-1658™) using the same protocol and ratio. Co-cultures and controls were evaluated at days 4 and 10 and subsequently analyzed by immunofluorescence staining.

### Co-culture of organoids with CAFs and PBMCs under shear stress in a flow-controlled bioreactor

PyMT-derived breast cancer organoids were dissociated and co-seeded with primary CAFs and autologous PBMCs (isolated from the same mouse). Organoids and CAFs were seeded at a 1:250 ratio into µ-Slide III 3D Perfusion chambers (Ibidi, 80371). Following seeding, chambers were incubated with breast cancer organoid expansion medium for 2 hours at 37 °C in a humidified atmosphere with 5% CO₂ to promote cell attachment. Chambers were then connected to a flow-controlled bioreactor system (57). The system included a peristaltic pump (ISMATEC MCP, 78002-00) equipped with an 8-roller, 12-channel pump head (ISMATEC Type IBM733A, 78002-36) and 1.52 mm PharMed BPT tubing (95723–36). µ-Slides were linked using Ibidi elbow luer connectors (10802) and serial connectors (10830) to sterile silastic tubing (Dow Corning 515-012). Organoid–CAF co-cultures were exposed to continuous capillary-level physiological shear stress (2.6 dyn/cm²). PBMCs were introduced into the circulating medium, allowing dynamic interaction with the organoid–CAF co-culture. The following experimental conditions were established: (1) Organoids alone under static conditions; (2) Organoids under flow (shear stress only); (3) Organoid–CAF co-culture under flow; (4) Organoid–CAF–PBMC co-culture under flow; (5) Organoid–CAF–PBMC co-culture under flow, treated with either THC or SR2. Cultures were analyzed at days 4 and 10 post-seeding.

### Mammary gland orthotopic injection in vivo

PyMT-derived organoids were treated with 10 nM THC for 4 days. After 10 days of cannabinoid withdrawal in culture, organoids were resuspended in basal medium and centrifuged at 500 × g. The resulting pellets were dissociated into single cells by incubating them for 20 minutes with TrypLE Select enzyme (Gibco, 12563011) at 37 °C with agitation. A total of 1 × 10⁴ organoid-derived cells in 100 μL (50 μL cell suspension and 50 μL BME) were orthotopically injected into the left inguinal mammary fat pad of 6-week-old PyMT wild-type sister mice, which served as controls for the organoid-derived injections. Tumor formation was first observed at week 3 post-inoculation for control (organoid-derived) tumors, and at week 5 for THC-pretreated organoid-derived tumors. Mice were monitored three times a week for tumor growth, with tumor length measured using external calipers. Tumor volume was calculated using the formula: (4π/3) x (width/2)^2^ x (length/2)^2^. When tumors reached a volume of 1000 mm³, mice were euthanized, and tumors were excised for evaluation. Tumors were fixed in 10% formalin and prepared for immunohistochemistry analysis.

### Analysis of metastasis in vivo

PyMT-derived organoids were differentiated using 10 nM THC for 4 days and subsequently cultured for an additional 14 days. Organoids were then minced and dissociated into a single-cell suspension by incubating with TrypLE for 20 minutes. A total of 5 × 10⁵ organoid-derived cells per mouse were injected into the lateral tail vein of 6-week-old PyMT wild-type mice to induce lung metastasis. Mice were euthanized four weeks post-injection. Lungs were then excised, fixed, and prepared for both immunohistochemistry and immunofluorescence analyses.

### Secondary organoid formation assay

Organoids were collected and dissociated into single cells to assess their self-renewal and organoid-forming potential. Samples were centrifuged at 500 × g, and the resulting pellets were incubated with pre-warmed TrypLE for 20 minutes at 37°C. Gentle mechanical dissociation was performed every 5 minutes. After dissociation, the samples were centrifuged again at 500 × g. Single cells were resuspended in BME at a density of 1 × 10⁴ cells per 40 μL and seeded into a 24-well plate. Following cell seeding, 650–1000 μL of BC organoid expansion medium was added to each well. Cells were maintained at 37°C in a humidified incubator with 5% CO₂. The medium was refreshed every two days, and organoid formation was evaluated after 10 days in culture.

### Cell cycle analysis

After completing the differentiation protocol, organoid cultures were harvested and centrifuged at 500 x g. Organoids were dissociated into single cells by incubating with TrypLE Select at 37°C for 20 minutes, with gentle pipetting every 5 minutes. The dissociated cells were then fixed in ice-cold 70% ethanol for 30 minutes on ice. After fixation, the cells were centrifuged at 800 x g. To degrade RNA and enable selective DNA staining, fixed cells were incubated with 100 μg/ml RNAse A (Sigma, R5503). Following RNA degradation, 400 μl of 50 μg/ml propidium iodide (Sigma, P-4170) was added per 1×10^6^ cells. The samples were incubated on ice for 10 minutes and then analyzed using a FACScalibur flow cytometer (BD Biosciences).

### Fluorescence-Activated Cell Sorting and subsequent organoid culture of PyMT:CB2R *tumors*

Mammary gland tumors from PyMT:CB2R^+/+^, and PyMT:CB2R^⁻/⁻^ mice were excised and processed for flow cytometry analysis. Tumors were minced and enzymatically digested in digestion solution (wash medium supplemented with 0.125 mg/mL Collagenase and 0.125 mg/mL Dispase II) for one hour at 37°C with shaking at 180 rpm. For single-cell suspension, the digested tissue was centrifuged at 600 × g and filtered through a 100 μm cell strainer using 10 mL of blocking buffer (wash buffer supplemented with 5 mM EDTA). The suspension was centrifuged again at 600 × g, and erythrocytes were lysed by incubating the sample for 15 minutes at room temperature. After a second filtration through a 100 μm cell strainer, the sample was centrifuged again at 600 × g and resuspended in FACS buffer (PBS supplemented with 2% FBS, 0.01% sodium azide, and antibiotics) at a concentration of 1 × 10⁶ cells per 100 μL. Fc receptors were blocked by incubating the cells with anti-CD16/32 antibody (1:100) for 20 minutes on ice. After blocking, cells were washed and resuspended at the same concentration for staining with the following antibodies: EpCAM-FITC (BioLegend, 118207) at a 1:200 dilution and CD49f-Alexa Fluor™ 647 (BioLegend, 313610) at a 1:20 dilution. Cells were incubated with the antibody mix for 20 minutes on ice in the dark. After staining, samples were centrifuged at 1000 × g at 4°C and resuspended in FACS buffer at a concentration of 1 × 10⁶ cells in 100 μL, and stained with 7-AAD (BioLegend, 420404) as a viability marker at a 1:50 dilution. Cells were sorted using a FACSAria III cell sorter (BD Biosciences) into four distinct populations based on differential expression of CD49f and EpCAM. FlowJo™ software was used for data analysis. Each population was collected separately and used to initiate organoid cultures. Sorted cells were seeded at a density of 1 × 10⁴ cells per well in 40 μL of BME in a pre-warmed 24-well plate. After matrix polymerization, 650– 1000 μL of BC organoid expansion medium was added to each well. Cultures were maintained at 37°C with 5% CO₂, with unsorted cells serving as the control for organoid formation. Medium was refreshed every 48 hours. Organoid formation was evaluated after 10 days in culture to assess the growth potential of each sorted population.

### Apoptosis assay by flow cytometry

Organoid cultures were harvested and transferred to a tube for dissociation into single cells. Pre-warmed TrypLE Select was used for cell dissociation, with incubation for 20 minutes at 37°C. After digestion, cells were centrifuged at 500 x g. Apoptosis analysis was performed using the Dead Cell Apoptosis Kit with Annexin V for Flow Cytometry (Invitrogen, V13445), according to the manufacturer’s instructions. Briefly, cells were stained with a 1:100 dilution of Annexin V-Alexa Fluor™ 488 and 0.2 μg/ml propidium iodide (PI) solution in annexin-binding buffer. The samples were incubated at room temperature for 15 minutes in the dark. Following incubation, samples were analyzed using a FACScalibur flow cytometer.

### Cell viability assay

Organoids and primary tumor cells derived from PyMT mice were assessed for viability based on intracellular ATP content using the CellTiter-Glo® 3D Cell Viability Assay (Promega, G9682), following the manufacturer’s instructions. Briefly, cells were lysed with the CellTiter-Glo 3D reagent, and luminescence was measured using an IVIS Spectrum system (PerkinElmer). Background signals from the medium were subtracted, and relative luminescence units (RLU) were normalized to control conditions.

### Cellular migration analysis from patient-derived samples

To investigate cellular migration traits in detail, we employed the HoloMonitor® M4, an automated live-cell imaging cytometer that performs time-lapse analysis. This system allows for the imaging and quantification of unstained, living cells directly within their culture vessels inside the incubator, enabling real-time monitoring of various morphological and migratory parameters. Based on digital holographic microscopy, the HoloMonitor measures phase shifts in light as it passes through cells, which the software then uses to reconstruct high-resolution topographic images. For our experiments, we used ibidi® plastic 24-well plates fitted with HoloLid™ lids— specifically designed to reduce image artifacts caused by surface vibrations and condensation. We quantified the following parameters: Cell area (μm²), the surface area occupied by each cell; Irregularity, the degree to which a cell’s outline deviates from a perfect circle; Migration (μm), the shortest linear distance between a cell’s initial and final positions; Motility (μm), the total path length traveled by a cell; Motility speed (μm/h), the motility distance divided by the observation time. Statistical analyses were performed using R software (version 4.5.0, RCore2013). For each parameter, we calculated the mean ± standard error of the mean (SEM). A generalized linear model (glm) (58) was fitted to assess the impact of experimental conditions, time and their interaction on each variable. Variables were log transformed to verify assumptions of normality and homoscedasticity.

### Cytokine profiling from co-culture conditioned media

Cytokine secretion profiles from co-culture conditions were analyzed using the Proteome Profiler Mouse Cytokine Array Kit, Panel A (R&D Systems, ARY006), according to the manufacturer’s protocol. Briefly, 1 mL of conditioned media was collected from cell cultures, centrifuged at 500 x g to remove cellular debris, and the supernatants were transferred to fresh tubes and stored at -20°C until processing. Array membranes were blocked and incubated with the Detection Antibody Cocktail. The antibody-sample mixture was applied to the membranes, which were incubated overnight at 4°C. The following day, membranes were washed, incubated with Streptavidin-HRP for 30 minutes, and developed using a chemiluminescent reagent. Signals were detected using the ChemiDoc XRS+ Imaging System (BioRad). Cytokine spot intensities were quantified using Fiji (ImageJ) and normalized to positive controls.

### Immunofluorescence

For immunofluorescence analysis, organoids were seeded into 8-well chamber slides with removable wells (Nunc™ Lab-Tek™ II Chamber Slide™ System 154534) during the final week of the culture period. Cells were briefly rinsed with phosphate-buffered saline (PBS) prior to fixation. Organoids were fixed with 10% formalin for 1 hour at room temperature (RT). Permeabilization was performed using 3% Triton X-100 (Sigma, T9284) in PBS for 20 minutes, followed by three washes with 0.1% Triton X-100 in PBS. Blocking was carried out with blocking solution (PBS + 0.1% Triton X-100 + 2% bovine serum albumin (BSA) [Sigma, A6003]) for 2.5 hours at RT. Primary antibodies were diluted in the blocking solution and added to the samples, which were then incubated overnight at 4°C. After incubation, samples were washed with PBS, and secondary antibodies were applied at a 1:500 dilution in the blocking solution. Incubation with secondary antibodies was done at RT for 1.5 hours in dark conditions. After secondary antibody incubation, samples were washed twice with PBS. For nuclear staining, DAPI was diluted 1:1000 in PBS and added to each well for a 10-minute incubation. After staining, the cells were mounted using Mowiol 4-88 (Calbiochem, 475904) as the mounting medium. The primary antibodies used for IF staining and their respective dilutions are as follows: **Cytokeratin 14** (rabbit monoclonal EPR17350, Abcam, ab181595) at 1:100 dilution. **Cytokeratin 8/18** (mouse monoclonal TROMA-I, DSHB, ab_531826) at 1:100 dilution. **Ki67** (mouse monoclonal B56, BD Biosciences, 550609) at 1:400 dilution. **E-cadherin** (rabbit monoclonal 24E10, Cell Signaling, 3195S) at 1:500 dilution. **Snail** (mouse monoclonal L70G2, Cell Signaling, 3895) at 1:250 dilution. **CD49f-Alexa Fluor™ 647** (rat monoclonal GoH3, BioLegend, 313610) at 1:100 dilution. **EpCAM** (rat monoclonal G8.8, BioLegend, 118207) at 1:200 dilution. **SOX10** (rabbit monoclonal EPR4007, Abcam, ab155279) at 1:250 dilution. **CD24-Alexa Fluor™ 488** (rat monoclonal M1/69, BioLegend, 101816) at 1:800 dilution. **H3K9me3** (rabbit polyclonal, Abcam, ab8898) at 1:400 dilution. **H3K4me3** (rabbit monoclonal EPR20551-225, Abcam, ab213224) at 1:100 dilution. **H3K27me3** (mouse monoclonal, Abcam ab6002) at 1:200 dilution. **Phalloidin-Rhodamine** (Invitrogen R415) at 1:200 dilution. Samples were examined using a TCS SP8 laser scanning confocal microscope (Leica). Image acquisition was performed in a sequential scanning mode by optimizing lasers and spectral detectors for each specific fluorescent dye. Z-stacks were obtained by 2.5 μm z-steps, a 20X oil immersion objective and a 1024×1024 scanning resolution. Image processing and analyses were performed using Fiji (ImageJ). 3D projections were generated using the brightest point projection method.

### Immunohistochemistry

Mammary gland and lung tissue samples were fixed in 10% formalin, paraffin-embedded, and sectioned at 4 μm. The sections were mounted on Superfrost Plus slides and dried overnight. Slides were stained with hematoxylin and eosin (H&E) or subjected to immunohistochemical analysis. Antigen retrieval was performed, followed by blocking of endogenous peroxidase activity using 3% hydrogen peroxide. The slides were then incubated with primary antibodies specific for the following targets: **Cytokeratin 14** (rabbit polyclonal AF64, Covance, PRB-155P), **Cytokeratin 8/18** (rabbit monoclonal, Dako, IR094), **SOX9** (rabbit polyclonal, Millipore, AB5535), **SOX10** (goat polyclonal N-20, Santa Cruz, c-17342), **Ki-67** (rabbit monoclonal D3B5, Cell Signaling, 12202), and **Estrogen Receptor alpha** (mouse monoclonal CRET94D/H9). Secondary antibodies, conjugated with horseradish peroxidase (OmniRabbit, Ventana, Roche), were applied, and the immunohistochemical reaction was developed using 3,3’-diaminobenzidine tetrahydrochloride (DAB) as the chromogen (Chromomap DAB, Ventana, Roche or DAB solution, Dako). Nuclei were counterstained with Carazzi’s hematoxylin. Finally, the slides were dehydrated, cleared, and mounted with a permanent mounting medium for microscopic evaluation.

### RNA extraction and analysis of mRNA levels

RNA was extracted using NucleoZOL™ (Takara Bio) following the manufacturer’s instructions. Reverse transcription (RT) was performed from 1 μg of RNA using the Transcriptor First Strand cDNA Synthesis Kit (Roche, 04379012001) with random hexamer primers. Quantitative real-time PCR was then conducted using the PowerUp™ SYBR™ Green Master Mix (Applied Biosystems, A25742), following the manufacturer’s guidelines, using 50 ng of cDNA per well in a 384-well plate. Data were acquired with a QuantStudio™ 12K Flex system (Applied Biosystems). The following primers were used (5′−3′ sequence; F: Forward, R: Reverse): **Krt5**. F: TCC TGT TGA ACG CCG CTG AC; R: CGG AAG GAC ACA CTG GAC TGG; **Krt14**. F: CAG CCC CTA CTT CAA GAC CA; R: GTC GAT CTG CAG GAG GAC AT; **Krt8**. F: GGA CAT CGA GAT CAC CAC CT; R: TGA AGC CAG GGC TAG TGA GT; **Krt18**. F: CAA GTC TGC CGA AAT CAG GGA C; R: TCC AAG TTG ATG TTC TGG TTT T; **Cdh1**. F: AGG AAA TGC ACC CCT CCA AT; R: AAT CGG CCA GCA TTT TCT G; **Cdh2**. F: GCC ATC ATC GCT ATC CTT CT; R: CCG TTT CAT CCA TAC CAC AAA; **Vimentin**. F: CCA ACC TTT TCT TCC CTG AAC; R: TTG AGT GGG TGT CAA CCA GA; **Sox10**. F: ATC AGC CAC GAG GTA ATG TCC AAC; R: ACT GCC CAG CCC GTA GCC; **EpCAM**. F: GAT TCT GCA CGT GAG ACC TG; R: GAT ACC AAG TCA AAC CGA GAA CTT.

### RNA sequencing

RNA was extracted using Nucleozol™ following the manufacturer’s instructions. RNA-seq sample preparation and initial data analysis were carried out by Novogene. In brief, transcriptome libraries were generated from high-quality RNA samples (triplicates), which were verified using the Agilent Bioanalyzer 2100. Messenger RNA was isolated using poly-T oligo-attached magnetic beads, then fragmented prior to cDNA synthesis. Both non-stranded and strand-specific libraries were prepared according to standard Illumina protocols. Library quality was assessed using Qubit, qPCR, and Bioanalyzer analysis. Sequencing was performed on an Illumina platform utilizing sequencing-by-synthesis technology to generate high-throughput reads. Bioinformatic analysis began with quality control of raw reads using FastQC. Clean reads were mapped to the reference genome with HISAT2. Differential expression was analyzed using DESeq2, applying a threshold of adjusted p ≤ 0.05 and |log2FC| ≥ 1. Functional enrichment was performed using the clusterProfiler R package, with analysis across GO, KEGG, Reactome, DO, and DisGeNET databases.

### Statistical analysis

All experiments were performed with at least three independent biological replicates. Data are presented as mean ± SEM, and p-values ≤ 0.05 were considered statistically significant. For comparisons between two independent groups, a two-tailed Student’s t-test was used. Correlations between two variables were calculated using Pearson’s r. For multi-group comparisons, data were analyzed using one-way ANOVA with post-hoc Tukey’s multiple comparison test, or two-way ANOVA when appropriate. Statistical analyses were performed using GraphPad Prism software (version 10.1.2).

## Data availability

RNAseq data has been deposited in the SRA repository under BioProject accession number PRJNA1276513.

## Acknowledgements

We thank the CNIO Histopathology and Genomics Units, UCM Confocal Unit and UCM Animal Facility for their technical support. We are indebted to the members of the Cannabinoid Signaling group (UCM), Eva Resel for administrative support, and to our former undergraduate and master students, with special mention to Marc Pinto, Manuel Luque, and Paula Gijón. We would like to express our thanks to Novogene Europe for their technical and customer support, with special appreciation to Metehan Cifdaloz, Paula Yagüe, and Haozhen Su. We are also extremely grateful to the Hospital *La Paz*, particularly to the Gynecology and Histopathology Units, for their constructive participation in this work and very specially, to the patients enrolled in this research study. This work has been in part financed by benefactors, through multiple donations to *Asociación Española contra el Cáncer* (AECC). We are extremely thankful to all our anonymous donors. The work has been funded by the Spanish Ministry of Science, Innovation and Universities, supported with European Regional Development funds: **CNS2022-135364** and **PID2022-136508OA-I00**, and the Complutense University, through the project **PR3/23-30841** to MS-R. MS-R was also supported by AECC (**INVES18005SALA**) and a *Ramón y Cajal* contract from the Ministry of Science, Innovation and Universities (**RYC2020-028929-I**). NGM-I was supported by AECC (**PRDMA19003GARC**) and Fulbright fellowship (**PS00380044**). ERE and MB were supported by National Institutes of Health (NIH) Research Project Grant **R01 HL 161069**.

## Author contributions

NGM-I conducted most of the cellular and *in vivo* experiments, with assistance from GP-G, MR-H, CQ-G, EM-Z, and MS-R. The Flow Cytometry and Confocal Imaging Unit at UCM (LMA-C and CH) collaborated on confocal imaging analyses and flow cytometry experiments. The Histopathology Unit at CNIO performed IHC experiments and provided professional histopathological analyses of the samples when needed (EC). MM-R and AZ-C contributed to RNA sequencing experiments, with support from NGM-I, CQ-G, and MS-R. AP-P collaborated with EM-Z and NGM-I to perform the R analysis of patient-derived data. LF-A assisted in preparing the necessary documents and protocols for patient enrollment at the hospital, recruited patients, and offered clinical advice. CS, MB-C and ERE provided guidance, materials and/or resources. MB-C and ERE directly contributed to the optimization and execution of dynamic cell co-culture experiments alongside NGM-I. MS-R and NGM-I supervised the project design, data evaluation and analysis. MS-R conceived the study, managed the teamwork and wrote the manuscript with contributions from the primary authors.

## Ethical approvals

All animal procedures were conducted at the Animal Facility of the Faculty of Biology, Complutense University of Madrid (UCM). The experimental protocols involving animals adhered to local regulations (RD53/2013 and ECC/566/2015) and were approved by the relevant Regulatory Agencies, including Complutense University and the Community of Madrid. All authors involved in animal experimentation hold the necessary accreditation (categories C and D) granted by the Competent Authority: CAP-T-0771-15, CAM Animal Welfare Committee on Animal Experimentation. Regarding patient protection, all procedures followed the guidelines of Hospital La Paz, local regulations, and the Declaration of Helsinki, as well as the Guidelines for Good Clinical Practice (GCP) established by the International Conference on Harmonization (ICH E6). The protocols for sample collection, processing, and any additional materials provided to the patients (e.g., Patient Information Sheets and the Informed Consent Document) were approved by the Clinical Research Ethics Committee of *Hospital La Paz*, in full compliance with national legislation.

