## Supplemental Figures for "Transient CB2 receptor activation triggers irreversible luminal differentiation via chromatin remodeling in breast cancer"

### Supplementary Figures

Supp. Figure 1. Short-term exposure to nanomolar THC induces long-term changes in the viability of MMTV-PyMT mammary tumor 3D organoids, but not in 2D cultures.

Supp. Figure 2. Brief exposure to nanomolar THC induces comparable effects in organoids derived from MMTV-PyMT and MMTV-Her-2/neu transgenic mammary tumors.

Supp. Figure 3. THC rewires differentiation programs in patient-derived breast cancer organoids.

Supp. Figure 4. Cannabis extracts derived from *Cannabis sativa* produce effects similar to those of pure phytocannabinoids on tumor cell differentiation, in contrast to endogenous cannabinoids.

Supp. Figure 5. Neither SR1 nor SR2 prevent the THC-mediated effects on tumor cell differentiation.

Supp. Figure 6. Knock-out of CB2R produces effects similar to those of THC on tumor cell differentiation.

Supp. Figure 7. Characterization of a 3D co-culture model integrating mammary tumor organoids, CAFs, and PBMCs under continuous shear stress.

Supp. Figure 8. Holographic imaging reveals reduced motility and impaired dispersal adaptation in SR2-treated organoid-derived cells during 2D migration.

### Supplementary figure 1

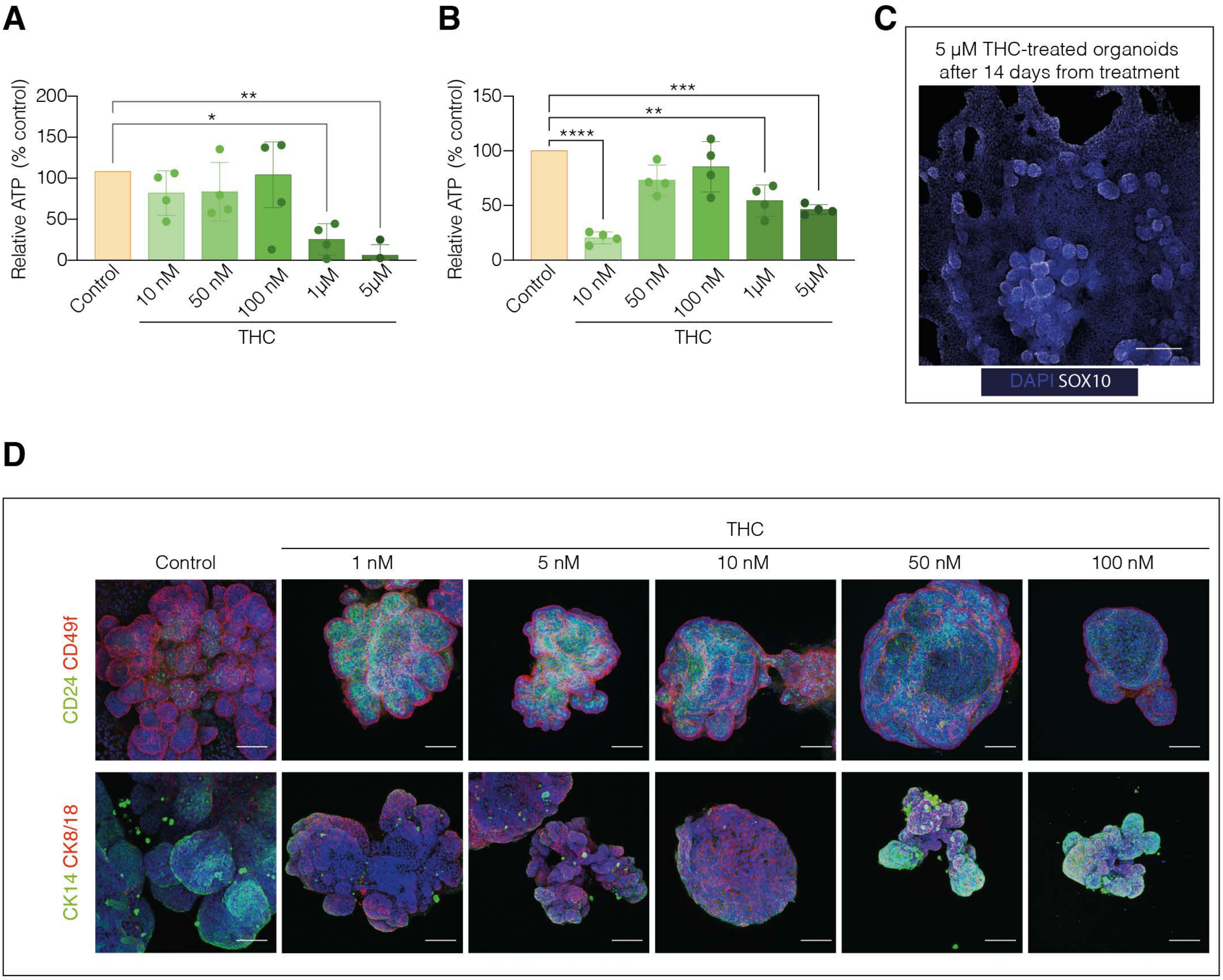

**Supp. Figure 1. Short-term exposure to nanomolar THC induces long-term changes in the viability of MMTV-PyMT mammary tumor 3D organoids, but not in 2D cultures. A,** Viability of 2D cell cultures derived from FVB-MMTV-PyMT mammary tumors, exposed to vehicle (DMSO) or varying THC concentrations for 4 days, with analysis 10 days post-treatment. *n*=3 independent experiments; *$p<0.05$; **$p<0.01$ (One-way ANOVA). **B**, Viability of 3D organoid cultures derived from FVB-MMTV-PyMT mammary tumors, treated similarly with vehicle or THC for 4 days, with analysis 10 days later. *n*=3 independent experiments; **$p<0.01$; ***$p< 0.001$; ****$p<0.0001$ (One-way ANOVA). **C**, Representative confocal images of THC-treated organoids (5 µM) assessed 14 days post-treatment. Organoids were stained for SOX10 (white) and DAPI for nuclei (blue). *n*=3 independent experiments. Scale bar: 100 µm. **D**, Representative confocal images of organoids treated with varying THC concentrations for 14 days, stained for CD24 (green) and CD49f (red) in the upper panels, or CK14 (green) and CK8/18 (red) in the lower panels. Note the changes in organoid shape and marker expression following 10 nM THC exposure. DAPI (blue) is used for nuclei staining. All images show 3D stacks of the examples depicted. *n*=3 independent experiments. Scale bar: 100 µm.

### Supplementary figure 2

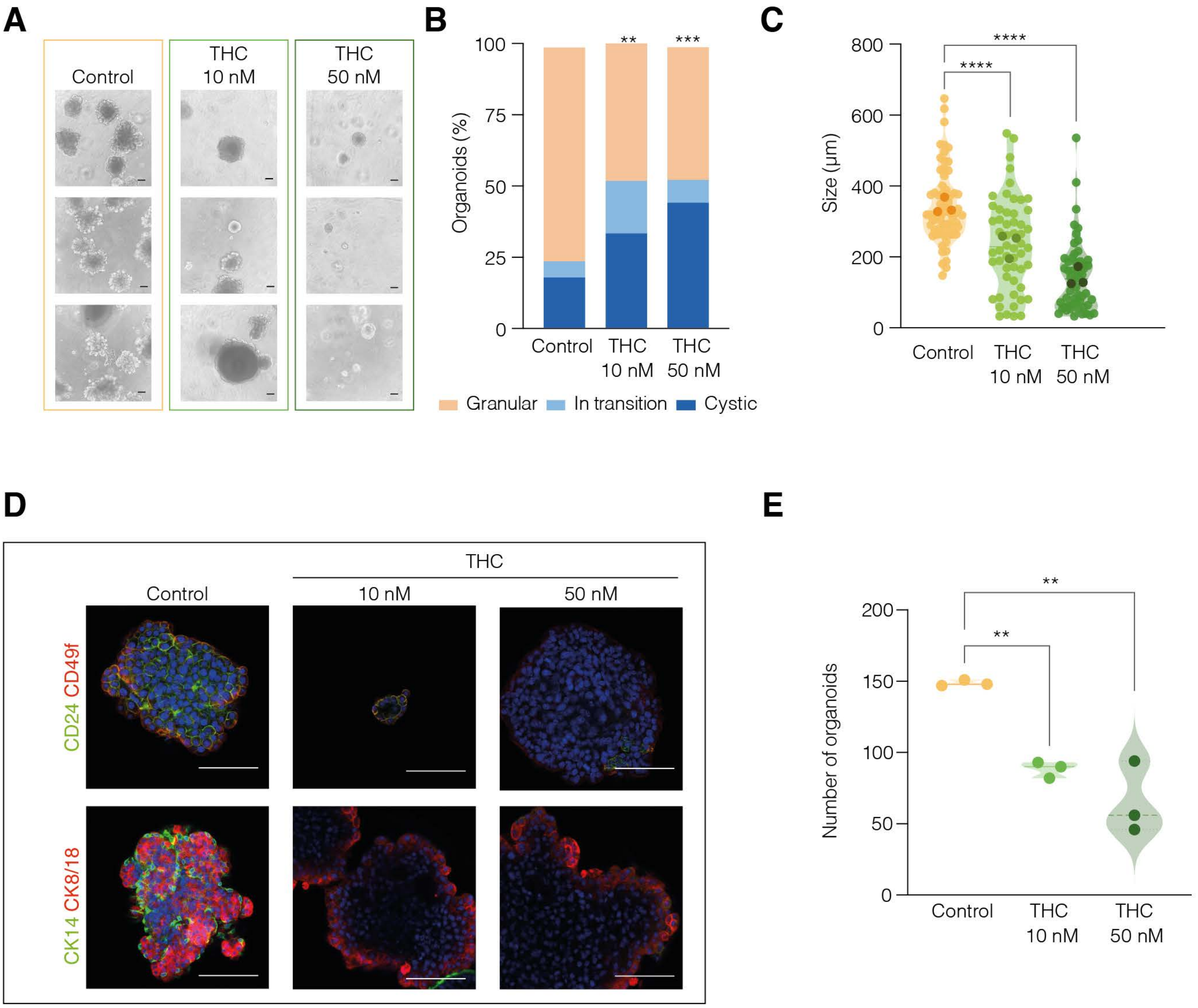

**Supp. Figure 2. Brief exposure to nanomolar THC induces comparable effects in organoids derived from MMTV-PyMT and MMTV-Her-2/neu transgenic mammary tumors.**

**A,** Representative brightfield images of MMTV-Her-2/neu-derived tumor organoids treated with vehicle (control), 10 nM THC, or 50 nM THC, following the schedule indicated in Fig. 1A. Organoids were analyzed at 2 weeks post-treatment. *n*=3 independent experiments. Scale bar: 100 μm. **B,** Quantification of organoid morphology at 14 days after treatment, categorizing them as granular (basal), mid (in transition), or cystic (luminal/rounded). Data show a significant increase in cystic organoids following THC treatment. *n*=3 independent experiments; **p<0.01; ***p<0.001 (Two-way ANOVA), comparing granular organoids vs. control. **C,** Quantification of organoid size (μm) for each treatment group, measured at 14 days post-treatment. THC treatment significantly reduced organoid size. *n*=3 independent experiments, with 80–100 organoids counted per condition; ****p<0.0001 (Two-way ANOVA). **D,** Confocal images of organoids treated with THC (10 nM or 50 nM) and stained for CD24 (green) and CD49f (red), or CK14 (green) and CK8/18 (red). Nuclei are stained with DAPI (blue). Images represent 3D stacks of organoids analyzed 14 days post-treatment. *n*=3 independent experiments. Scale bar: 100 μm. **E,** Total number of secondary organoids generated from single-cell dissociation, indicating the self-renewal capacity of the cultures, evaluated 21 days after treatment. THC-treated cultures exhibited significantly lower self-renewal compared to controls. *n*=3 independent experiments; **p<0.01 (Two-way ANOVA).

### Supplementary figure 3

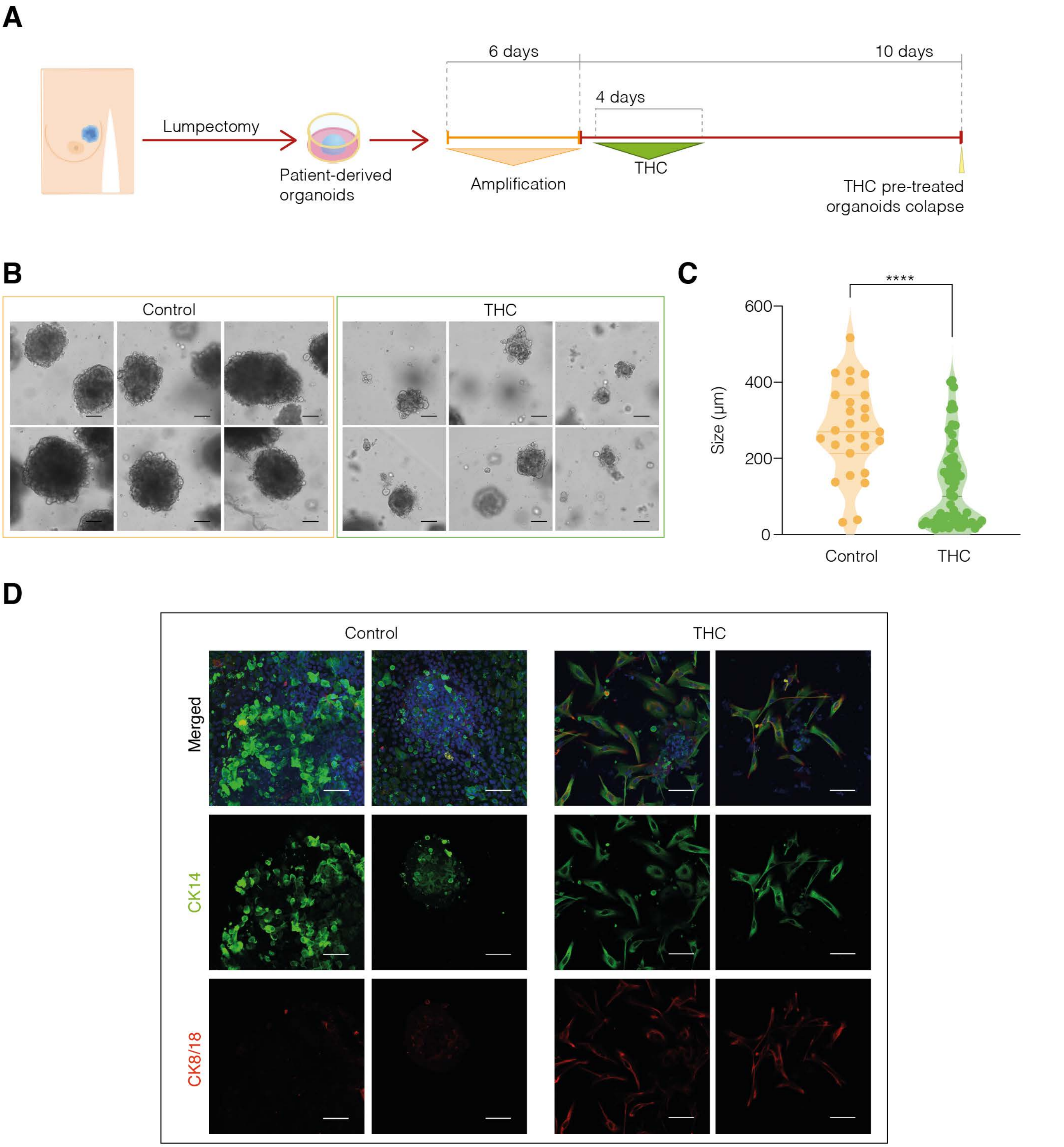

**Supp. Figure 3. THC rewires differentiation programs in patient-derived breast cancer organoids. A**, Experimental design. Surgery samples were used to establish patient-derived organoid cultures. After amplification, organoids were treated with vehicle or THC (10 nM) for 4 days and maintained for one week without treatment. Cannabinoid exposure significantly reduced self-renewal capacity, leading to culture collapse by ~day 10. **B**, Representative brightfield images of organoids treated with vehicle or THC (10 nM), analyzed one week after treatment. *n*=3 independent experiments. Scale bar, 100 µm. **C**, Quantification of organoid size (µm) in both conditions. *n*=3; 80–100 organoids per condition; ****$p<0.0001$ (unpaired Student's *t*-test). **D**, Confocal images of organoids stained for CK14 (green) and CK8/18 (red), analyzed one-week post-treatment. DAPI stains nuclei (blue). Shown are 3D stacks of representative examples. *n*=3; scale bar, 100 µm.

### Supplementary figure 4

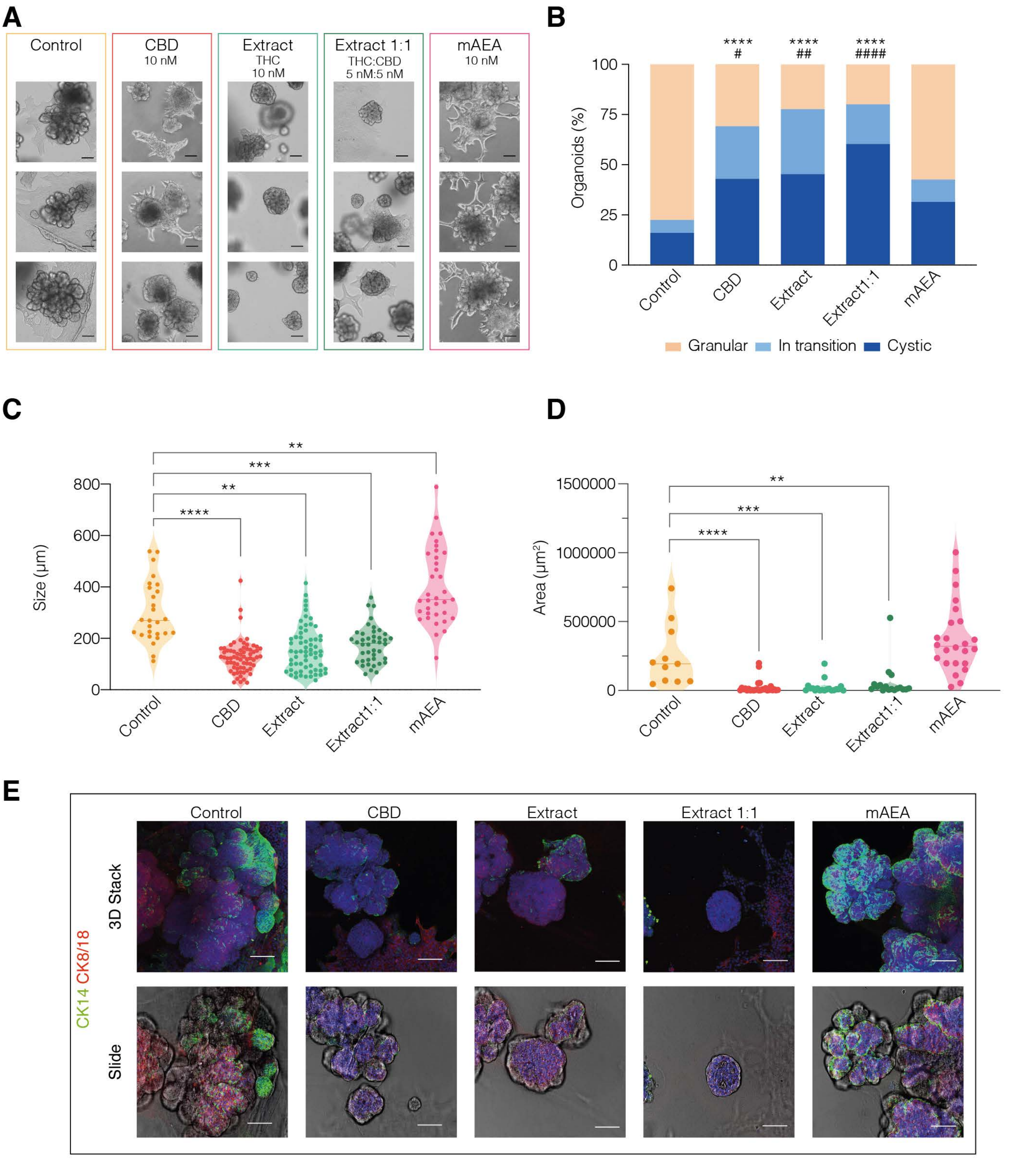

**Supp. Figure 4. Cannabis extracts derived from *Cannabis sativa* produce effects similar to those of pure phytocannabinoids on tumor cell differentiation, in contrast to endogenous cannabinoids. A,** Representative brightfield images of organoid cultures treated with vehicle, CBD (10 nM), THC-rich extract from *Cannabis sativa* (10 nM, "Extract THC"), a THC:CBD 1:1 extract (5 nM each, "Extract 1:1 THC:CBD"), or methanandamide (mAEA, a stable anandamide analogue, as a representative endogenous CB2 receptor ligand). Organoids were treated for 4 days, followed by 2 weeks of drug withdrawal, and analyzed at the end of the period. *n*=3 independent experiments. Scale bar: 100 μm. **B,** Quantification of organoid morphology at 14 days post-treatment, categorizing them as granular (basal), mid (in transition), or cystic (luminal/rounded). THC and CBD-rich extracts significantly increased the cystic morphology compared to controls. *n*=3 independent experiments; ****p<0.0001 (Two-way ANOVA), comparing granular organoids vs. control; $^{\#}$p<0.05; $^{\#\#}$p<0.01; $^{\#\#\#\#}$p<0.0001 (Two-way ANOVA), comparing cystic organoids vs. control. **C,** Quantification of organoid size (μm) for each treatment condition, measured 14 days post-treatment. Significant size reduction was observed in organoids treated with THC or CBD-rich extracts. *n*=3 independent experiments, with 80–100 organoids counted per condition; **p<0.01; ***p<0.001; ****p<0.0001 (One-way ANOVA). **D,** Quantification of the two-dimensional area (μm²) of collective-migratory projections in organoids treated with the five conditions, evaluated at 14 days. THC and CBD-rich extracts significantly reduced migratory projections compared to control. *n*=3 independent experiments, with 25–30 organoids counted per condition; **p<0.01; ***p<0.001; ****p<0.0001 (unpaired Student's t-test). **E,** Representative confocal images of organoids treated as indicated and stained for CK14 (green) and CK8/18 (red) at 14 days post-treatment. Nuclei are stained with DAPI (blue). Upper panels show 3D stacks, and lower panels show representative slices. *n*=4 independent experiments. Scale bar: 100 μm.

### Supplementary figure 5

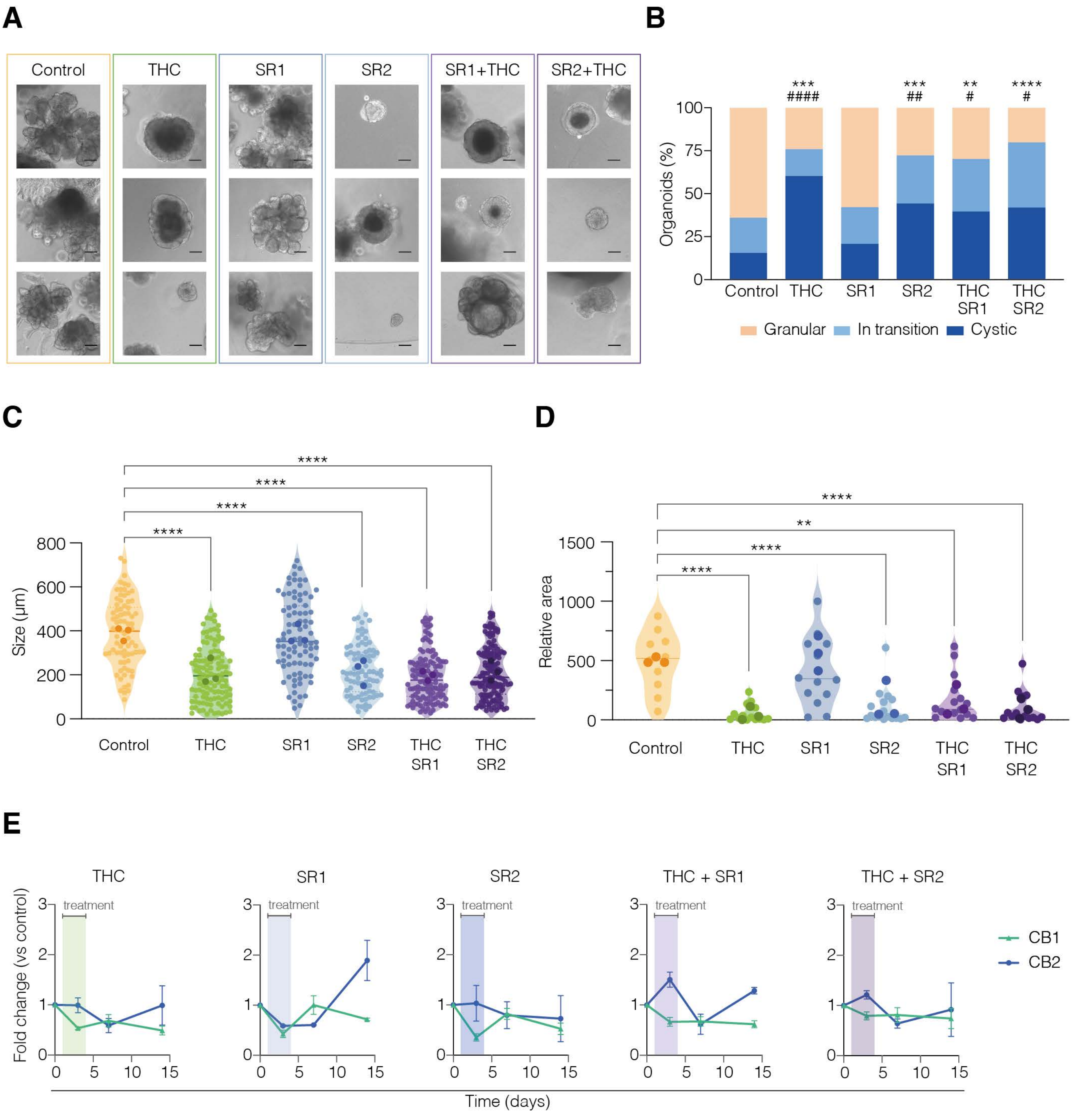

**Supp. Figure 5. Neither SR1 nor SR2 prevent the THC-mediated effects on tumor cell differentiation. A,** Representative brightfield images of organoid cultures treated with vehicle (DMSO), THC (10 nM), CB1R inverse agonist SR1 (100 nM), CB2R inverse agonist SR2 (100 nM), or combinations of THC with SR1 or SR2. Treatments were applied for 4 days, followed by 2 weeks of drug withdrawal. *n*=3 independent experiments. Scale bar: 100 µm. **B,** Quantification of organoid morphology at 14 days post-treatment, categorizing them as granular (basal), mid (in transition), or cystic (luminal/rounded). No significant differences were observed between THC alone and the combinations with SR1 or SR2. *n*=3 independent experiments; **p<0.01; ***p<0.001; ****p<0.0001 (Two-way ANOVA), comparing granular organoids vs. control; $^{\#}$p<0.05; $^{\#\#}$p<0.01; $^{\#\#\#\#}$p<0.0001 (Two-way ANOVA), comparing cystic organoids vs. control. **C,** Quantification of organoid size (µm) in all conditions at 14 days post-treatment. No significant changes were observed between THC and the combinations with SR1 or SR2. *n*=3 independent experiments, with 80–100 organoids counted per condition; ****p<0.0001 (One-way ANOVA). **D,** Quantification of the two-dimensional area (µm$^2$) of collective-migratory projections in organoids treated with the five conditions, analyzed at 14 days. No significant difference was found between THC and the combinations with SR1 or SR2. *n*=3 independent experiments, with 10–20 organoids counted per condition; **p<0.01; ****p<0.0001 (unpaired Student's t-test). **E,** mRNA expression levels of CB1R (green) and CB2R (blue) measured by quantitative PCR at various time points: before, during, or after treatment. mRNA levels were normalized to GAPDH housekeeping genes. *n*=3 independent experiments; no significant changes detected (unpaired Student's t-test).

### Supplementary figure 6

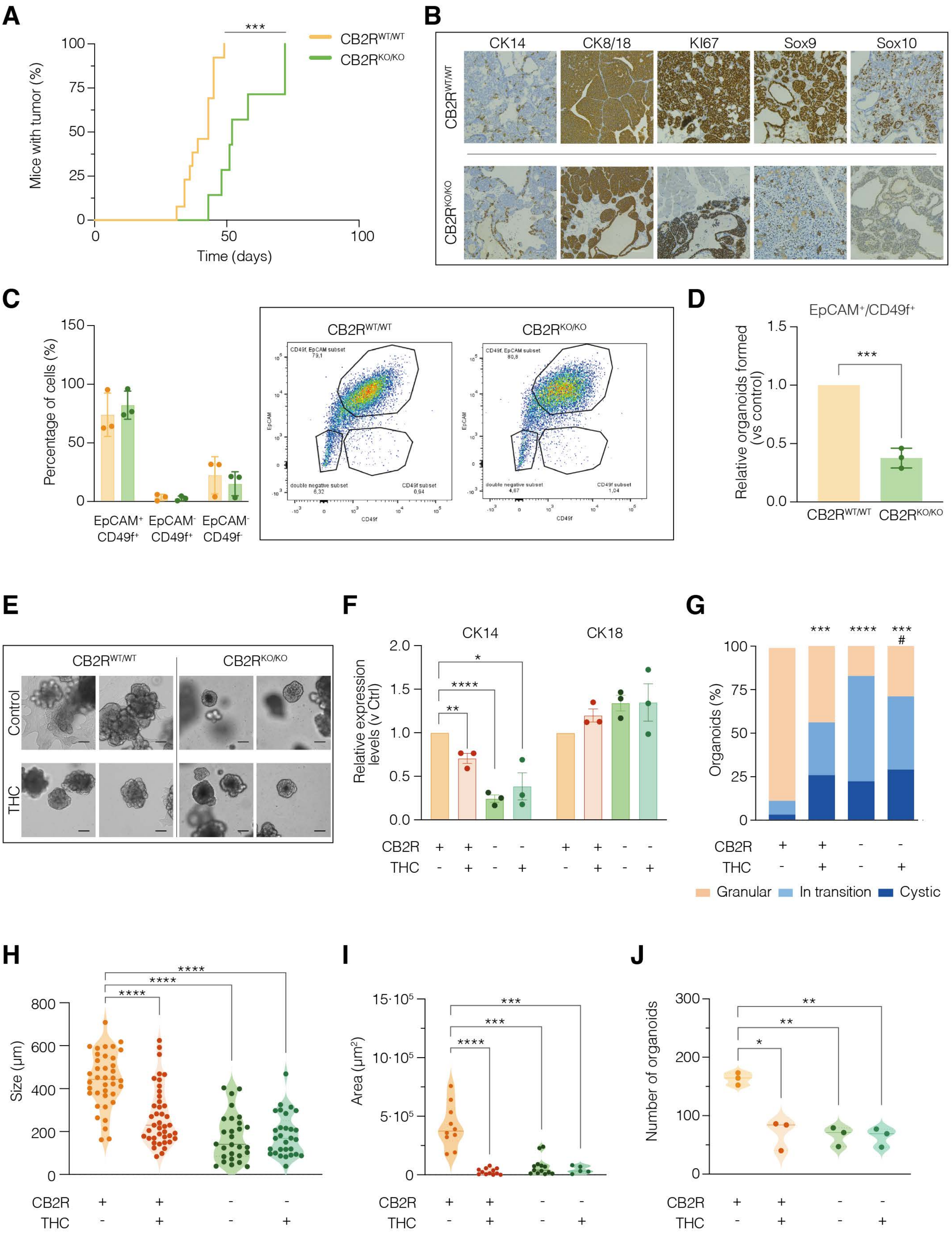

**Supp. Figure 6. Knock-out of CB2R produces effects similar to those of THC on tumor cell differentiation. A,** Kaplan-Meier plot showing tumor onset in MMTV-PyMT CB2R$^{WT/WT}$ and CB2R$^{KO/KO}$ mice over 100 days, measured as the percentage of mice with tumors. CB2R$^{WT/WT}$ *n*=15, CB2R$^{KO/KO}$ n=12 animals; ***p<0.001 (Two-way ANOVA). **B,** Representative images of tumor molecular characterization. Protein expression levels of CK14 (basal marker), CK8/18 (luminal marker), KI67 (proliferation marker), SOX9 and SOX10 (stemness markersl were evaluated by immunohistochemistry. Scale bar: 500 µm. **C,** Flow cytometry analysis of EpCAM/CD49f subpopulations in CB2R$^{WT/WT}$ and CB2R$^{KO/KO}$ tumors: EpCAM+/CD49f+ (luminal progenitors), EpCAM-/CD49f+ (stem-like mammary cells), EpCAM-/CD49f- (differentiated basal cells). Left panel shows the percentage of cells in each subpopulation. Right panel shows representative flow cytometry plots for each subpopulation. *n*=3 independent experiments. **D,** Relative comparison of the number of organoids generated from the EpCAM+/CD49f+ subpopulation (luminal progenitors) of tumor cells. The data indicates impaired stemness potential in luminal progenitors from CB2R$^{KO/KO}$ tumors compared to CB2R$^{WT/WT}$ tumors. Only organoids derived from luminal progenitors are shown, as other EpCAM/CD49f subpopulations did not generate stable organoid cultures. *n*=3 independent experiments; ***p<0.001 (unpaired Student's t-test). **E,** Representative brightfield images of organoid cultures derived from CB2R$^{WT/WT}$ and CB2R$^{KO/KO}$ tumors treated with vehicle or THC (10 nM). Treatments were applied for 4 days, followed by 2 weeks of drug withdrawal. *n*=3 independent experiments. Scale bar: 100 µm. **F,** mRNA expression levels of basal (CK14) and luminal (CK18) markers, measured by quantitative PCR. mRNA levels were normalized to GAPDH. *n*=3 independent experiments; *p<0.05; **p<0.01; ****p<0.0001 (unpaired Student's t-test). **G,** Quantification of organoid morphology at 14 days post-treatment, categorizing them as granular (basal), mid (in transition), or cystic (luminal/rounded). *n*=3 independent experiments; ***p<0.001; ****p<0.0001 (Two-way ANOVA), comparing granular organoids vs. control; $^{\#}$p<0.05 (Two-way ANOVA), comparing cystic organoids vs. control. **H,** Quantification of organoid size (µm) at 14 days post-treatment. *n*=3 independent experiments, with 80–100 organoids counted per condition; ****p<0.0001 (One-way ANOVA). **I,** Quantification of the two-dimensional area (µm²) of collective-migratory projections in organoids analyzed at 14 days. *n*=3 independent experiments, with 10–20 organoids counted per condition; ***p<0.001; ****p<0.0001 (unpaired

Student's t-test). **J,** Total number of secondary organoids generated from single-cell disaggregation (reflecting self-renewal capacity) in the four conditions tested, evaluated at 21 days post-treatment. *n*=3 independent experiments; *$p<0.05$; **$p<0.01$ (One-way ANOVA).

### Supplementary figure 7

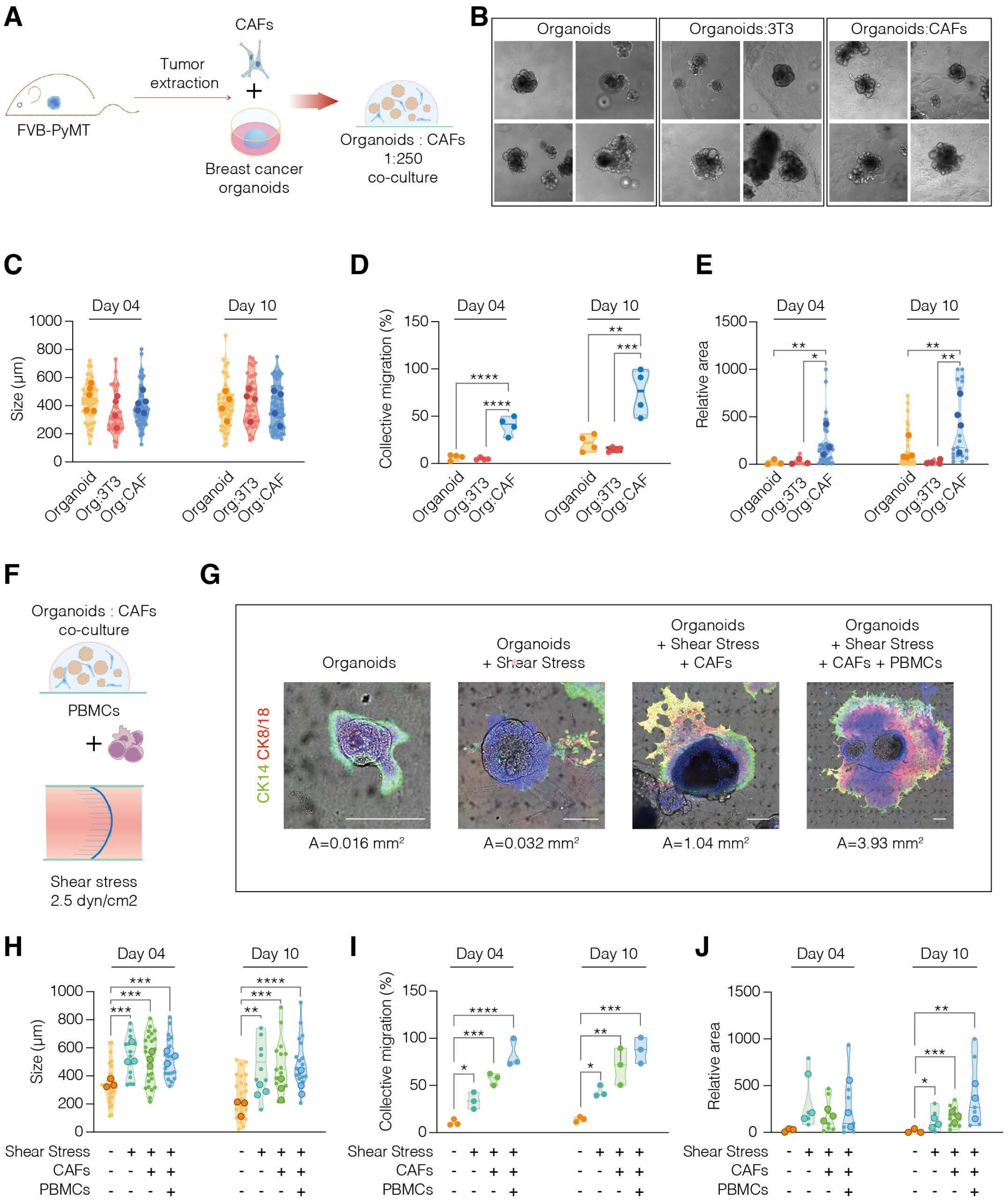

**Supp. Figure 7. Characterization of a 3D co-culture model integrating mammary tumor organoids, CAFs, and PBMCs under continuous shear stress**. **A**, Schematic of the initial co-culture setup. MMTV-PyMT–derived organoids were embedded in BME and co-cultured with either primary cancer-associated fibroblasts (CAFs) or NIH3T3 fibroblasts at a 1:250 ratio. **B**, Representative brightfield images of organoids cultured alone (left), with 3T3 fibroblasts (middle), or with CAFs (right) at day 10. **C**, Quantification of organoid size (μm) at day 4 and day 10 across the conditions shown in (B). *n*=4 independent experiments; ns, not significant (One-way ANOVA). **D**, Percentage of organoids displaying collective migration at day 4 and day 10. *n*=4 independent experiments; **p<0.01; ***p<0.001; ****p<0.0001 (One-way ANOVA). **E**, Quantification of collective migratory projection area (μm²) across conditions at day 4 and day 10. *n*=3 independent experiments; *p<0.05; **p<0.01 (One-way ANOVA). **F**, Schematic of the advanced 3D co-culture model. Organoids were embedded in BME, co-cultured with syngeneic CAFs (1:250), and exposed to circulating peripheral blood mononuclear cells (PBMCs) under continuous shear stress (2.6 dyn/cm²) via pumped flow. **G**, Confocal images of organoids stained for CK8/18 (red) and CK14 (green), with DAPI (blue) marking nuclei, under the conditions shown in (F) at day 10. Representative slices of collective migration are included. Quantified area (A) corresponds to the region of collective migration. *n*=3 independent experiments. Scale bar, 250 μm. **H**, Organoid size (μm) measured at day 4 and day 10 in the conditions shown in (F). *n*=3 independent experiments; **p<0.01; ***p<0.001; ****p<0.0001 (One-way ANOVA). **I**, Percentage of organoids with collective migration at day 4 and day 10. *n*=3 independent experiments; *p<0.05; **p<0.01; ***p<0.001; ****p<0.0001 (One-way ANOVA). **J**, Quantification of collective migratory projection area (μm²) at day 4 and day 10. *n*=3 independent experiments; *p<0.05; **p<0.01; ***p<0.001 (Unpaired Student's t-test).

### Supplementary figure 8

**A**

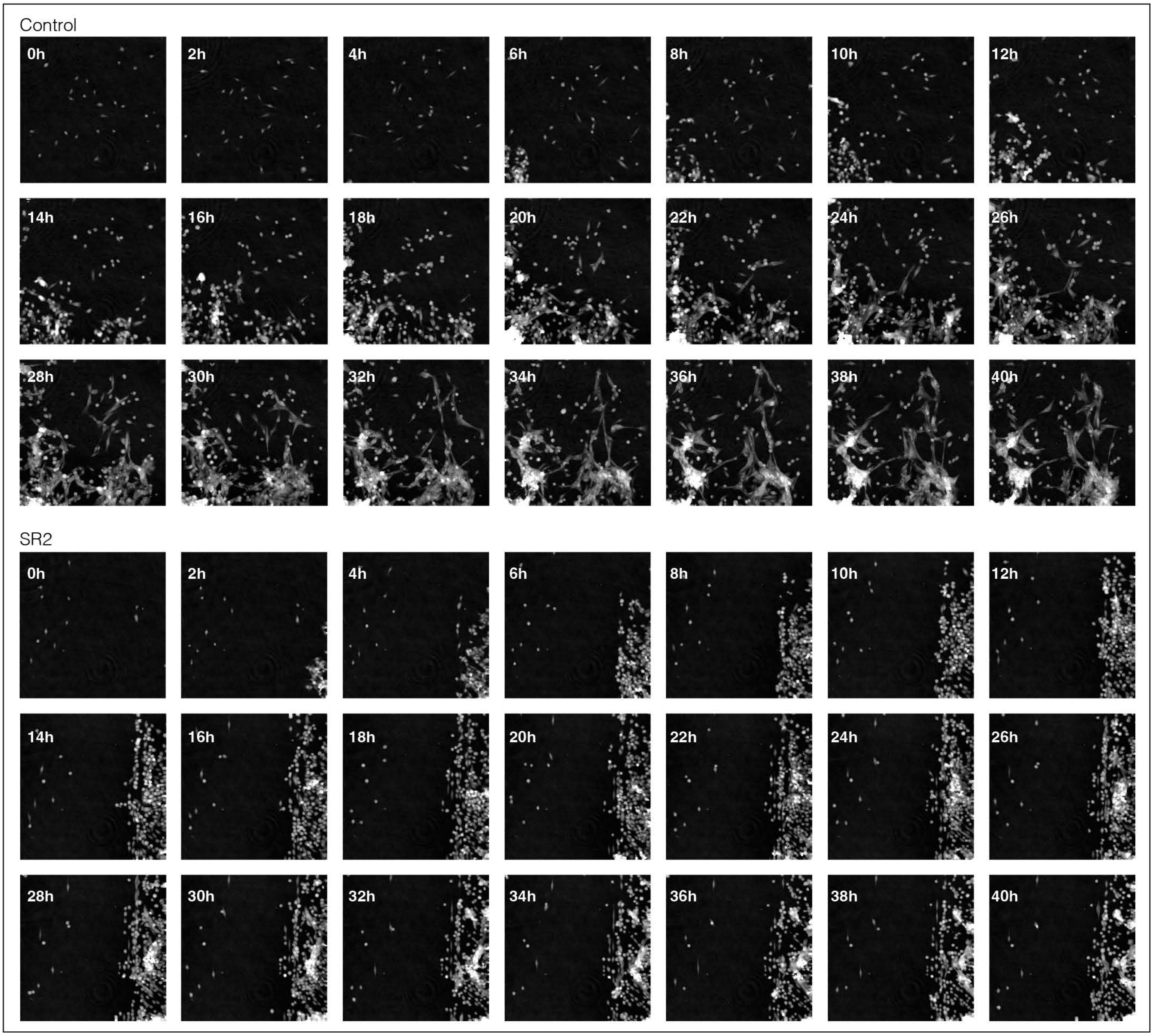


**B**

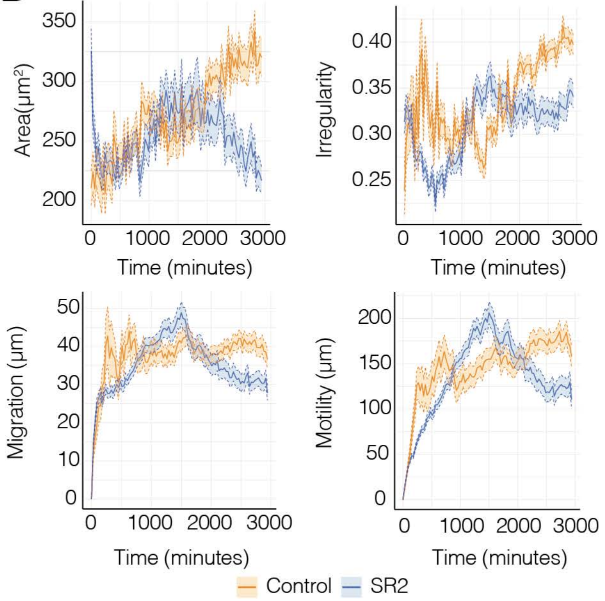


**C**

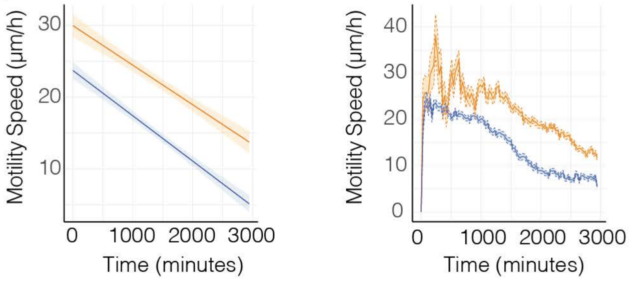


**D**

| | Group*time p-value | | Group p-value | |
|---|---|---|---|---|
| Area | <2e$^{-16}$ | **** | 1.29 e$^{-08}$ | **** |
| Irregularity | <2e$^{-16}$ | **** | 0.00288 | ** |
| Migration | 0.0381 | * | 0.0228 | * |
| Motility | 0.0684 | | 0.0740 | |
| Motility speed | 0.851 | | <2e$^{-16}$ | **** |

**Supp. Figure 8. Holographic imaging reveals reduced motility and impaired dispersal adaptation in SR2-treated organoid-derived cells during 2D migration. A**, Phase holographic images showing 2D migration of patient-derived organoid cells treated with vehicle or SR2 (100 nM), assessed via wound healing assay at time points from 0 h to 40 h. **B**, Quantification of key wound healing parameters—cell area ($\mu m^2$), shape irregularity, migration distance (µm), and motility (µm)—under both treatment conditions. Data are presented as mean ± standard error of the mean (SEM). **C**, Quantification of migration speed (µm/h) under treatment conditions. Mean ± SEM (left panel) and a fitted linear model to evaluate the effect of the treatment (right) are shown. **D**, Statistical analyses performed in R (version 4.5.0) of the indicated parameters. For all cases, model residuals were inspected to confirm assumptions of normality and homoscedasticity.
